# Single-cell transcriptomic resolution of stem cells and their developmental trajectories in the hippocampus reveals epigenetic control of cell state perseverance

**DOI:** 10.1101/2021.07.21.452775

**Authors:** Adrián Salas-Bastos, Martin Treppner, Josip S. Herman, Dimitrios Koutsogiannis, Harald Binder, Michael B. Stadler, Dominic Grün, Tanja Vogel

**Affiliations:** Institute of Anatomy and Cell Biology, Department of Molecular Embryology, Medical Faculty, Albert-Ludwigs-University Freiburg, 79104 Freiburg, Germany; Faculty of Biology, Albert-Ludwigs-University Freiburg, Freiburg, Germany; Institute of Medical Biometry and Statistics, Faculty of Medicine and Medical Center, Albert-Ludwigs-University Freiburg, 79104 Freiburg, Germany; Freiburg Center for Data Analysis and Modeling, Albert-Ludwigs-University Freiburg, 79104 Freiburg, Germany; Max Planck Institute of Immunobiology and Epigenetics, Freiburg, Germany; International Max Planck Research School for Molecular and Cellular Biology (IMPRS-MCB), Freiburg, Germany; Würzburg Institute of Systems Immunology, Julius-Maximilians-University Würzburg, 97078 Würzburg, Germany; Friedrich Miescher Institute for Biomedical Research, Basel, Switzerland; SIB Swiss Institute of Bioinformatics, Basel, Switzerland; Faculty of Science, University of Basel, Basel, Switzerland; Centre for Integrative Biological Signaling Studies, Albert-Ludwigs-University Freiburg, 79104 Freiburg, Germany; Center for Basics in NeuroModulation (NeuroModul Basics), Medical Faculty, Albert-Ludwigs-University Freiburg, 79104 Freiburg, Germany; Freiburg Institute for Advanced Studies (FRIAS), Albert-Ludwigs-University Freiburg, 79104 Freiburg, Germany

## Abstract

Despite conceptual research on hippocampus development and the application of single-cell-resolved technologies, the nature and maturation of its diverse progenitor populations are unexplored. The chromatin modifier DOT1L balances progenitor proliferation and differentiation, and conditional loss-of-function mice featured impaired hippocampus development. We applied single-cell RNA sequencing on DOT1L-mutant mice and explored cell trajectories in the E16.5 hippocampus. We resolved in our data five distinct neural stem cell populations with the developmental repertoire to specifically generate the cornu ammonis (CA) 1 field and the dentate gyrus (DG). Within the two developing CA1- and CA3-fields, we identified two distinct maturation states and we thus propose CA1- and CA3-differentiation along the radial axis. In the developing hippocampus, DOT1L is primarily involved in the proper development of CA3 and the DG, and it serves as a state-preserving epigenetic factor that orchestrates the expression of several important transcription factors that impact neuronal differentiation and maturation.

**Graphical Abstract:** 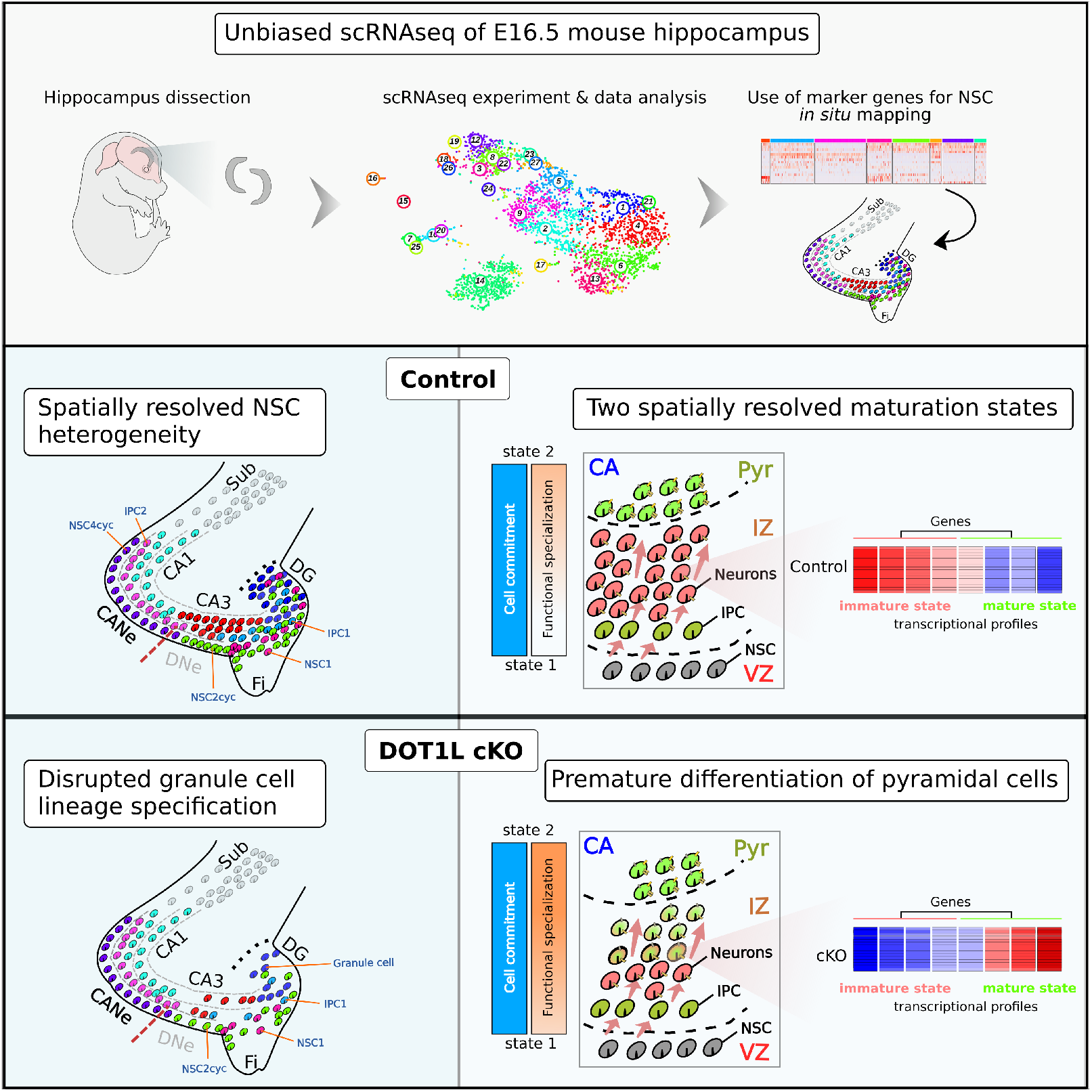

- The developing hippocampus contains distinct and spatially separated NSC populations that differ in expression of a specific set of firstly described marker genes.
- CA pyramidal neurons mature along the radial axis and pass through distinct maturation states.
- DOT1L preserves the dentate granule cell lineage in the developing hippocampus and limits maturation in the CA1- and CA3-fields development.
- DOT1L gates cell maturation as upstream regulator of transcription factor expression that confer instrumental roles in hippocampus development.

## Introduction

The hippocampus develops from the medial pallium of the dorsal telencephalon^1^. As part of the limbic system, the hippocampal neuronal networks are important for learning, memory, emotions and other functions^2^. It contains three neuronal layers, and is a conserved structure among vertebrates^3^. But surprisingly, up to date, the hippocampus, that is heavily studied to understand such specific function as spatial navigation and orientation, is poorly understood in its developmental origins.

The mature hippocampus is composed of the cornu ammonis (CA) fields and the dentate gyrus (DG). The development of both regions starts in embryonic and continues during postnatal stages^4–6^. CA and DG neurons originate from neural stem cells (NSCs) that locate to the ventricular zone (VZ)^4, 7^ and generate glutamatergic pyramidal cells (PC) or granule cells (GC). Cell migration of committed hippocampal progenitors and differentiating neurons along the radial and longitudinal axes represent crucial processes during embryonic development of the CA-fields^8, 9^ and DG^10^.

Several studies report on transcription factors (TFs) impacting hippocampus development^11–21^ and they give valuable insights into how this process is controlled transcriptionally. It is still unclear, but highly likely, that upstream signals coordinate the expression of instructing TFs, which probably act in organised networks. Epigenetic mechanisms that converge on concerted enhancer activation or on histone modifications to balance activation/repression of TF expression in a spatio-temporal manner, could be in place to orchestrate transcriptional programs driven by individual TFs. We here present the histone methyltransferase *disruptor-of-telomeric-silencing-1-like*, DOT1L, as upstream regulator of TFs that highly impacts hippocampus development by balancing progenitor proliferation and differentiation.

DOT1L conditional mouse mutants (cKO) present a disrupted cytoarchitecture of the hippocampus in which the DG is seemingly lost^22^. DOT1L impacts development of various parts of the central nervous system (CNS)^23, 24^. In all brain regions studied, DOT1L deficiency accelerates neuronal differentiation and migration. Thus, the alterations in the hippocampus in DOT1L-cKO suggested that studying the loss-of-function (LOF) of DOT1L will provide novel insights into general mechanisms of neuronal differentiation during hippocampus development. Using single-cell transcriptome (scRNAseq) analysis of control and mutant E16.5 hippocampus we resolved spatially distinct progenitor populations exerting region-specific transcriptional programs. We reconstructed neuronal differentiation along the radial dimension and different maturation trajectories which were characterised by expression of a distinct set of marker genes. We showed that DOT1L preserved hippocampus identity and repelled expansion of the subiculum. DOT1L prevented premature differentiation of precursors and maturation of committed neurons through concerted transcriptional activation of key TFs. We provide an integrated high-resolution view of the lineage trajectories in place during hippocampus development, identify the central genetic regulators and establish that the histone modifier DOT1L acts as state-preserving epigenetic factor preventing untimely neuronal differentiation and maturation.

## Results

### DOT1L deficiency changes proportions of NSCs, IPCs, CA3 PCs and DG GCs

Emx1-cre;Dot1l^fl/fl^ mice (DOT1L-cKO) presented with a disorganised CA and DG region (Figure 1A), similar to the phenotype observed using Foxg1-cre (**Figure S1A**). We performed scRNAseq^25^ on E16.5 hippocampus from one control and two DOT1L-cKO (Figure 1B), using 4135 cells. This sample size was estimated to be suitable for subsequent clustering analyses^26^ (**Figure S2**). 3701 cells remained after pre-processing, and RaceID^27^ analysis resolved 27 clusters (Figure 1C), most of which were annotated using known marker genes^6, 10, 15, 28–30^ (Figure 1D). All expected cell-types were identified, including five clusters of NSCs. We observed a separation of CA1 from CA3 PCs and dentate GCs, and that the latter two related transcriptionally closer to each other than to CA1 (Figure 1D). We did not observe any major influence of the sex nor the genotype on the cell distribution (**Figure S3**).

**Figure 1.**
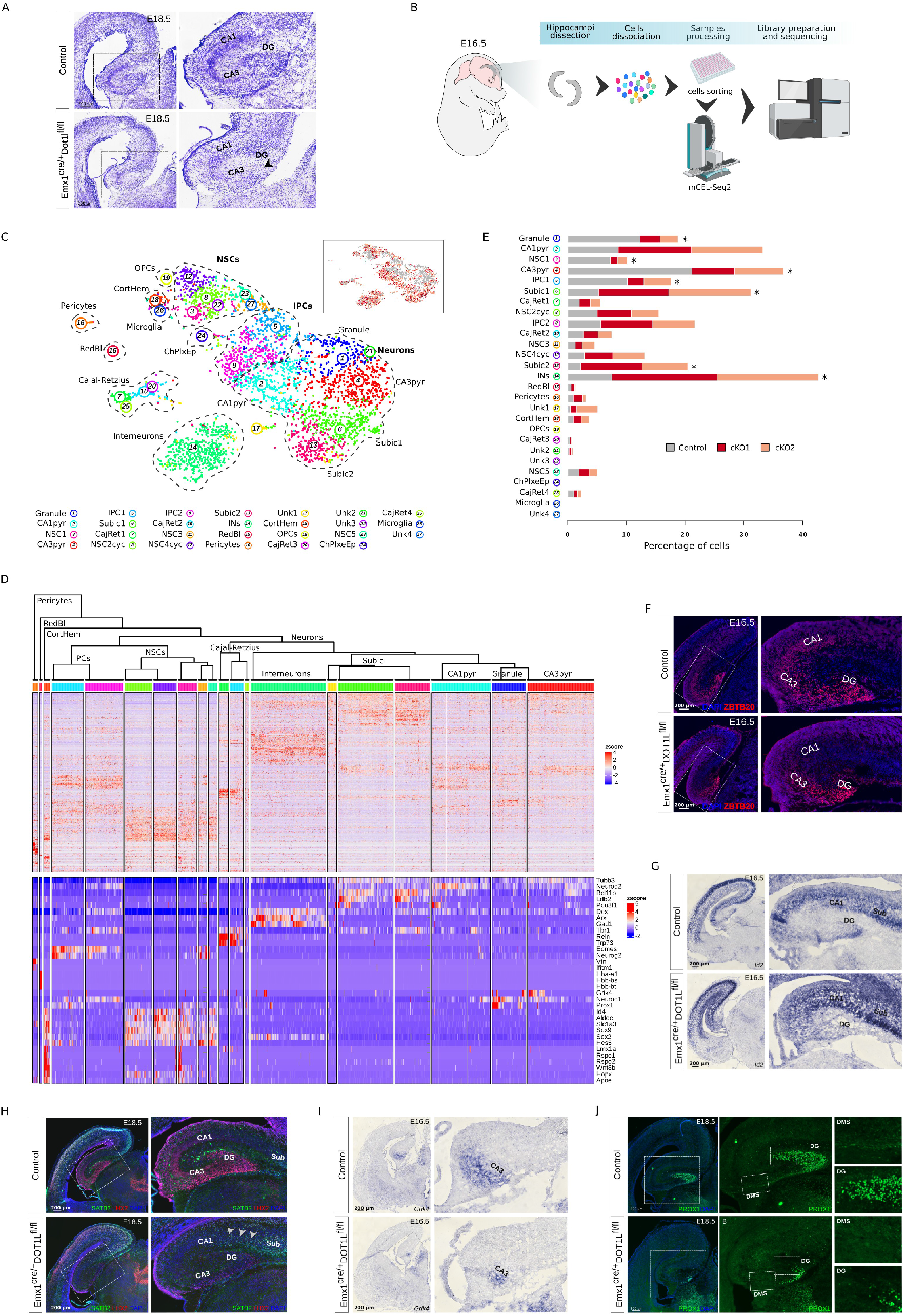
DOT1L-cKO alters hippocampal cytoarchitecture by affecting specific cell populations. A) Nissl staining on E18.5 control and cKO brain sections (n = 3), showing apparent loss of the DG region in the mutants (black arrowhead). B) Experimental design. E16.5 hippocampi of control and cKO littermates were dissected, followed by cell dissociation, single cells FACS sorting and libraries preparation using the mCEL-Seq2 protocol. C) tSNE plot showing the 27 clusters obtained after RaceID analysis. Inset shows the distribution of control (grey) and cKO (red and salmon) cells in the tSNE representation. D) Heatmaps showing the z-score normalized expression (red: high; blue: low) of highly variable genes (top) and exemplary marker genes (bottom) used for cell cluster annotation. Only clusters containing at least 10 cells are shown in the heatmaps. Hierarchical clustering of cell populations was performed on highly variable genes based on euclidean distance. E) Barplot showing the cell proportions in each cluster normalized by sample. Clusters significantly changing in proportions based on Fisher’s exact test (adjusted P<0.05) are indicated with asterisks. F) Immunofluorescence staining for ZBTB20 on E16.5 control and mutant brain sections showing depletion of positive cells in the cKOs (n = 3) in different areas of the hippocampal region. G) *In situ* hybridization for *Id2* on E16.5 control and cKO brain sections indicating high expression at the level of the putative CA1 in the mutants. H) Immunofluorescence co-staining for LHX2 and SATB2 on E18.5 control and mutant brain sections showing depletion of LHX2 positive cells in the cKOs (n = 3) in different areas of the hippocampal region and expansion of the SATB2 positive cells over the CA1 domain (white arrowheads). I) *In situ* hybridization for *Grik4* on E16.5 control and cKO brain sections (n = 3). J) Immunofluorescence staining for PROX1 on E18.5 control and mutant brain sections (n = 3) showing depletion of positive cells in the cKOs in the DG region and along the DMS. Scale bars are indicated inside each respective image. Sub: Subiculum; CA: cornu ammonis; DMS: dentate migratory stream; DG: dentate gyrus; Pyr: pyramidal cell layer; IZ: intermediate zone.

Although all clusters contained both control and mutant cells (Figure 1C**, inset**), an in-depth analysis of the distribution of cells per cluster for each animal showed a significant decrease of the proportion of DG granule, NSC1, CA3 pyramidal and IPC1 clusters upon DOT1L-LOF (adjusted P<0.05; Fisher’s test) (Figure 1E). The subiculum (Subic1, Subic2) and interneuron (INs) clusters showed significantly higher cell proportions for each of the two DOT1L-cKO samples compared to controls (adjusted P<0.05; Fisher’s test) (Figure 1E). Thus, DOT1L-cKO quantitatively altered the cell-type composition in the developing hippocampus in a cell-type specific manner.

We studied the enrichment of DOT1L-cKO cells with expression of subiculum marker genes and the boundary to the adjacent CA1-field. ZBTB20-expressing cells localized in the intermediate zone (IZ), the CA PC layer and the DG, but they were largely absent in the subiculum (Figure 1F). The ZBTB20 expression domain in DOT1L-cKO seemed smaller and localised ventro-laterally with fewer positive cells in the DG region compared to controls (Figures 1F**, S4A**). The proportions of CA1 PCs did not decrease significantly in the scRNAseq data suggesting a spatial re-arrangement of CA1 and subiculum, whereby the subiculum was shifted ventro-laterally upon DOT1L-LOF. ISH for *Id2* as cortical/subiculum marker^20^ showed an expanded subiculum expression domain reaching the prospective CA1-field ventrally in DOT1L*-*cKOs (Figure 1G). SATB2-positive cells, present in the subiculum but mostly absent from the CA1-field^31^, also extended ventrally in DOT1L-cKOs (Figure 1H). LHX2-positive cells, marking the hippocampal/cortical boundary^13^, were almost completely absent in the DOT1L-cKOs in the hippocampal VZ, CA-fields and DG, compared to their presence in controls (Figure 1H). Foxg1-cre DOT1L*-*cKO (**Figures S1B-D**) displayed similar alterations, which strongly indicated that DOT1L preserved hippocampal identity.

ISH for *Grik4*^6^ confirmed the decreased representation of CA3 PCs upon DOT1L-LOF compared to controls (Figure 1I). Further, we confirmed the impaired DG in the DOT1L-cKO, because we observed fewer PROX1- and EOMES-positive cells compared to controls along the dentate migratory stream (DMS) and in the developing DG in DOT1L-cKOs (Figures 1J, 4B**, S1E,F)**. PAX6-expressing GC progenitors of the DG presented a similar pattern as PROX1-positive cells in the DOT1L-cKOs (**Figure S4C**), underlining reduced presence of DG granule precursors and neurons upon DOT1L-LOF. Summarising, DOT1L prevented expansion of subiculum identity into the hippocampus and regulated cell numbers that settle in the CA3-field and DG.

### Lineage trajectory analyses resolve different progenitor populations in the developing hippocampus

Studies of hippocampus development incompletely resolved transcriptional programs underlying spatio-temporal cell fate trajectories of progenitors, but rely on broadly expressed NSC and IPC markers, including *Pax6*, *Sox9, Eomes* or Notch signalling pathway members^10^. Due to the high resolution inherent to the scRNAseq technology we resolved five different NSC and two IPC clusters (Figures 1C,D) in the E16.5 hippocampus. RNA velocity analysis^32^ using scvelo^33^ resolved cell trajectories (Figure 2A), in which NSCs clustered together and tended to move towards IPC1, followed by IPC2. IPC2 could thus represent a mature intermediate state towards neuronal differentiation compared to IPC1. IPC2 connected to the CA1 PCs, followed by CA3 PCs and DG GCs. Notably, the velocity vectors for the subiculum clusters converged with both CA PC populations, but they lacked direct connection either to NSCs or IPCs identified in our scRNAseq data set (Figure 2A).

**Figure 2.**
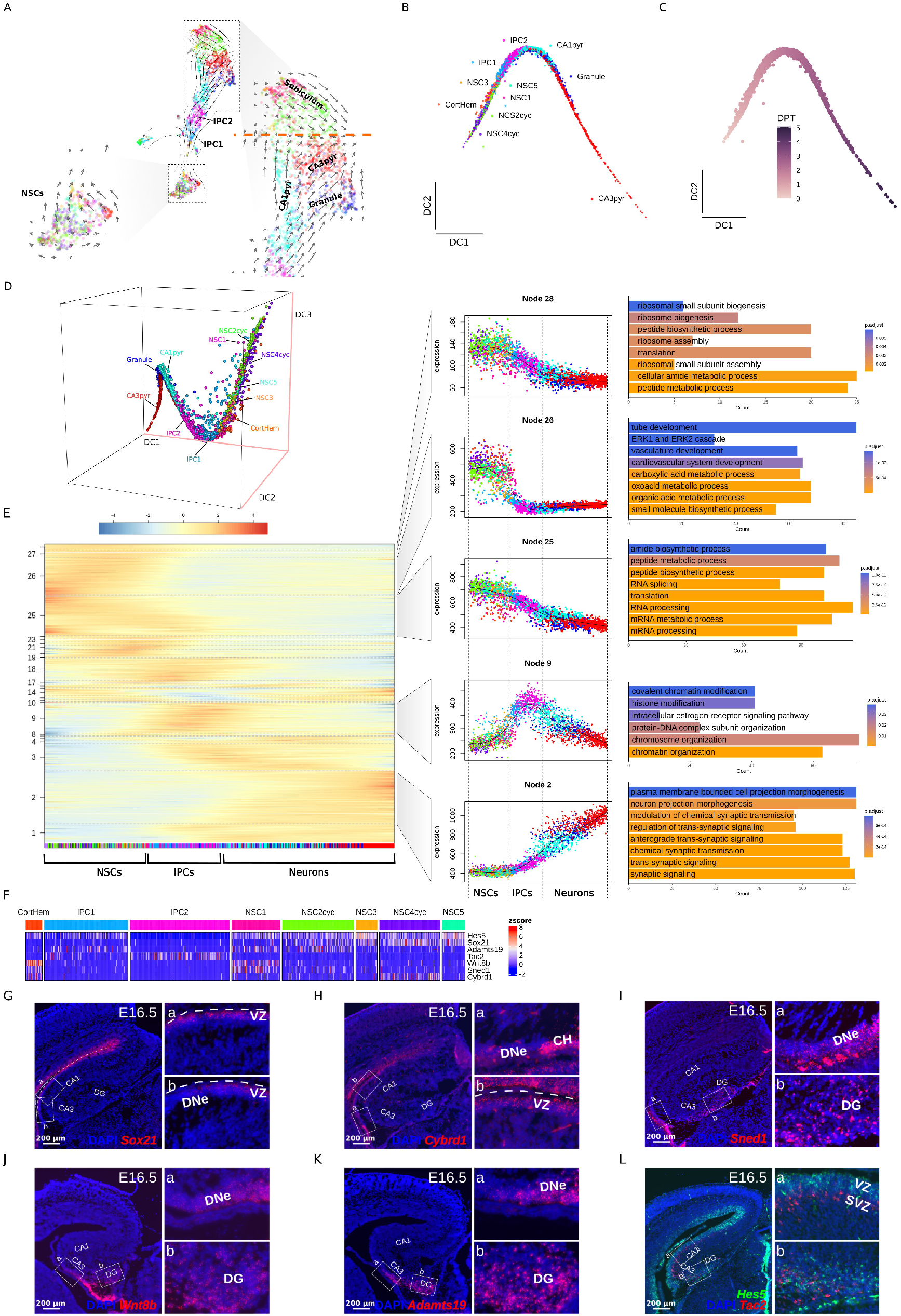
Lineage trajectory analysis identifies distinct and spatially separated progenitor populations in the developing hippocampus. A) scvelo stream plot showing averaged velocity vectors (arrows) indicative of the main transitions between the different cell clusters identified at E16.5. Magnification panels for the NSCs and committed neuronal populations are shown. Red dotted line highlights the point where the adjacent subiculum and CA pyramidal cell populations meet each other. Clusters are color coded as in Figure 1C. B) Two-dimensional diffusion maps plot showing the lineage trajectory of the hippocampal populations. Clusters are color coded as in Figure 1C. C) Two-dimensional diffusion pseudotime plot showing increase in the pseudotime as cells progress from the NSC to the committed cell states. D) Three-dimensional diffusion maps plot depicting the separation of NSC4cyc and NSC2cyc populations, apparently intermingled in B. Clusters are color coded as in Figure 1C. Red lines in the axes indicate the 3 distinct diffusion components (DC). E) Self organizing map showing the expression levels of gene sets (nodes) changing along the diffusion pseudotime axis (left). The running mean expression for selected nodes is shown for the different cell clusters (middle). Barplots indicating the main biological processes enriched (*q-value* < 0.05) in each highlighted node are shown (right). F) Heatmap showing expression levels for putative marker genes for the NSCs and IPCs clusters. G-L) smFISH stainings for selected NSCs and IPCs markers showing their spatial expression domains on E16.5 control brain sections (n = 3). Scale bars are indicated inside each image panel. DC: diffusion component; CA: cornu ammonis; DNe: dentate neuroepithelium; DG: dentate gyrus; VZ: ventricular zone; SVZ: subventricular zone; CH: cortical hem.

To resolve the connections between the different NSC clusters in detail, we applied diffusion maps (DM)^34^ and diffusion pseudotime (DPT)^35^ analyses, after filtering out cells unrelated to the hippocampal lineage (see methods). DM and DPT analyses showed an increased pseudotime as cells moved from the NSC (left) towards the more committed cell populations (right). CA3 PCs appeared as the most mature population at E16.5 (Figures 2B,C). NSC2cyc and NSC4cyc localized closest to the trajectory start. Due to transcriptional similarities within stem cells, they seemingly intermingled with each other and with the remaining NSC1,3,5 and cortical hem (CH, CortHem) clusters (Figure 2B). The cell arrangement based on the first three diffusion components indicated that NSCs conserved partly their cluster structure (Figure 2D), with differences between the respective NSC populations. DMs corroborated sequential development from the NSCs to the IPC1 and IPC2 states, followed by CA1 PCs, DG GC and CA3 PCs (Figures 2 B,D). We investigated the main gene sets and corresponding biological processes that changed along the trajectory by combining the cell order provided by the DPT with self-organizing maps (Figure 2E). Our analysis retrieved a series of nodes representing groups of genes highly expressed at different stages along the pseudotime axis. NSCs expressed strongly genes related to metabolic processes (Figure 2E, Node 26), and both NSCs and IPCs expressed gene sets associated with translation and mRNA processing (Figure 2E, Nodes 25 and 28). Genes included in these nodes decreased in expression with advanced differentiation. As cells matured towards IPCs and neuronal states, gene sets involved in synaptic function increased in transcription (Figure 2E, Node 2). Noticeable, genes affecting chromatin organization and histone modifications increased in expression from NSCs to IPCs, had a maximum expression in IPCs, and decreased as differentiation moved ahead on the diffusion pseudotime axis (Figure 2E, Node 9). IPC1 preceded IPC2 cells in this node, corroborating different maturation states in the intermediate progenitors.

DM analysis indicated seemingly hierarchical ordered NSC populations, but could not delineate a clear distinction of the different NSC clusters. We thus defined, based on visual inspection of the expression pattern of the DEGs between populations (adjusted P<0.05) in the tSNE representation, marker genes of the respective NSC and IPC subtypes and used single-molecule FISH (smFISH) to resolve the spatial locations of the distinct cell populations (Figure 2F). *Sox21* transcription delineated the CH, NSC1 and NSC2cyc (low *Sox21*) from NSC3, 4cyc, and 5 (high *Sox21*). *Sox21-*signals were strongest in the cortical VZ and CA neuroepithelium (CANe), not extending into the dentate neuroepithelium (DNe), origin of the DG (Figure 2G). *Cybrd1* expression marked NSC4cyc and CH, and smFISH showed that NSC4cyc cells located basally in the CANe, compared to apically located *Sox21*-expressing cells, which confined NSC3 and 5 within the VZ of the CANe. Consistent with its expression in CH cells, a defined *Cybrd1*-expressing cell population localized near the fimbria (Figure 2H).

*Sned1, Wnt8b* and *Adamts19* expression characterised NSC1 (expression of all three markers, *Sned1* higher) and NSC2cyc (*Adamts19* high; *Sned1*, *Wnt8b* low) (Figures 2I-K). According to *Sned1* expression, NSC1 localised in the VZ of the DNe extending towards the DG. NSC2cyc localised to the same area. Although NSC1 and NSC2cyc had similar expression domains, expression of *Sox21* in the NSC2cyc (Figure 2F) probably localises these cells apically in the DNe. scRNAseq and smFISH together resolved two spatially separated progenitor regions: NSC3, NSC4cyc and NSC5 precursors comprise the CANe, whereas NSC1 and NSC2cyc demarcated the DNe.

The lineage trajectories suggested that IPC1 and 2 are common, probably consecutive states during differentiation, originating from both progenitor populations (CANe, DNe). We selected marker genes for discriminating both IPC populations and characterizing their spatial location: all NSCs, CH and IPC1 expressed strongly *Hes5*, but had low expression in IPC2; *Tac2* was almost exclusively expressed in IPC2 with the exception of some few IPC1 (Figure 2F). *Hes5/Tac2* co-staining showed that *Hes5-*expressing cells mainly localized along the entire VZ of the CANe and DNe (Figure 2L). *Tac2-*expressing cells localized in the SVZ along the CANe and DNe, extending into the DG (Figure 2L). The expression domain of both markers seemed well separated between VZ and SVZ in the CANe but positive cells appeared intermingled in the DG region. IPC1 showed higher levels of *Adamts19* compared to IPC2 (Figure 2F), and *Adamts19* expression was restricted in the DNe and DG (Figure 2K). Thus, IPC2 are basal progenitors in the SVZ of the CANe, and *Hes5/Adamts19*-positive IPC1 localised to the DNe and DG. Together with the bioinformatics prediction that IPC1 seemingly preceded IPC2, we interpreted that IPC1 and IPC2 are two spatially separated progenitor fractions that were seemingly depicted in an earlier (IPC1) and advanced (IPC2) maturation state. Their distribution reflected probably the slightly advanced development of the CA1 PC compared to DG GC.

### NSCs and IPCs are biased towards CA1 PC and DG GC fate at E16.5

Our data suggested two separate germinative regions to generate the CA-fields and DG. To investigate the connections between the specific NSC/IPC populations to the mature cell lineages we performed CellRank^36^ and FateID^37^ analyses. As we knew the cell populations that represented the final cell states, we provided this information to the algorithms (Figure 3A). CellRank (Figure 3B) and FateID (Figure 3C) results highlighted distinct fractions of the NSCs with high probabilities to generate either CA1 PC or DG GC. Specifically, FateID analysis highlighted that CA1 PC fate connected to the NSC3, NSC4cyc, and NSC5 clusters (Figure 3C). This observation was in line with the NSC spatial expression domains we identified *in vivo* (Figures 2G-H) and confirmed the separation of the CANe and DNe. Both algorithms indicated that IPC1 had a fate bias towards DG GC, whereas IPC2 seemed to generate CA1 PCs at E16.5. This observation supported that IPC2 connected to the advanced development of CA1 as suggested by their localisation *in vivo*. The different NSC and IPC populations identified at E16.5, however, had very low fate probabilities towards the CA3 PC and subiculum cell lineages. CellRank analysis additionally indicated a gradually declining differentiation potential, which was highest in the NSCs, moderate in IPC, and lowest in neurons. Within the IPCs, IPC2 had a lower differentiation potential compared to IPC1, confirming that the former represented a slightly advanced intermediate state (Figure 3B).

**Figure 3.**
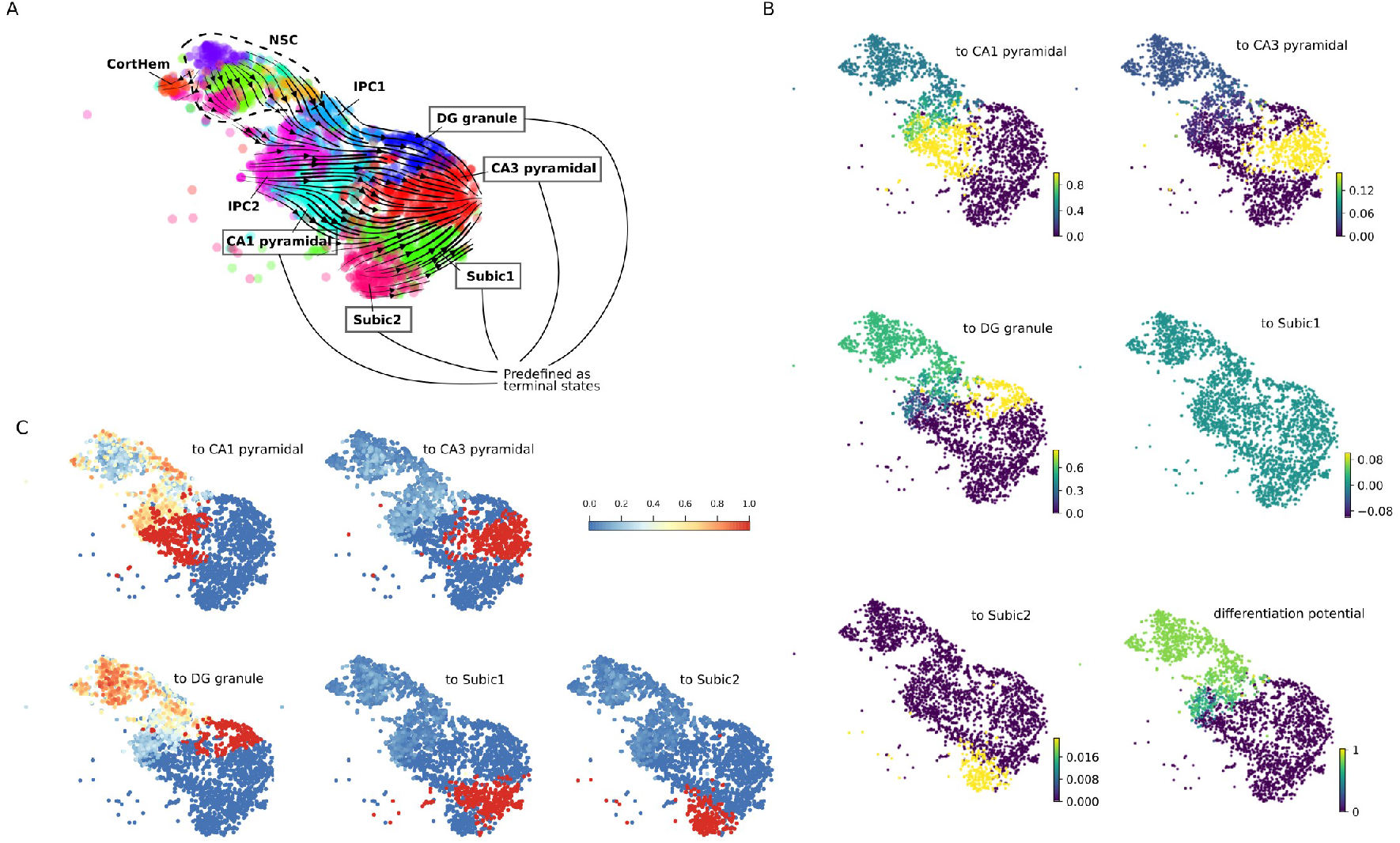
NSCs and IPCs show cell fate biases towards CA1 pyramidal or DG granule cells in the E16.5 hippocampus. A) Stream plot showing the average velocity vectors projected on the tSNE representation of the hippocampal populations. The mature cell populations preset as terminal states are indicated in the figure. Clusters are colour coded as in Figure 1C. B) tSNE plots showing the differentiation potential and the estimated fate probabilities towards the distinct terminal states as computed by CellRank analysis. Scale bars indicate the probability values. C) tSNE plots showing the estimated fate probabilities towards the distinct terminal states as computed by FateID analysis. Scale bar indicate the probability values.

### DOT1L preserves the DNe and prevents premature differentiation

We used the newly identified marker gene expressions in the stem cell compartment of the developing hippocampus to study by smFISH the specific NSCs/IPCs affected in DOT1L-cKOs. *Cybrd1* expression in the CANe (NSC4cyc) appeared similar in both genotypes (Figure 4A). *Sned1*-positive NSC1 within the DNe/DG region reduced largely upon DOT1L-cKO (Figure 4B). DNe/DG-located *Hes5*-expressing IPC1 decreased strongly in the DOT1L-cKOs compared to the CANe, where *Hes5*-expression was unaffected (Figure 4C). *Tac2*-positive IPC2 in the SVZ were kept in the DOT1L-cKO CANe, but reduced in the DNe/DG. These observations confirmed strong reduction of the stem cell populations residing in the DNe/DG (NSC1, IPC1) upon DOT1L-cKO, and seemingly normal CANe appearance. The finding of fewer DG stem cells explained reduced numbers of PROX1-positive DG GC in DOT1L-cKO (Figure 1J).

**Figure 4.**
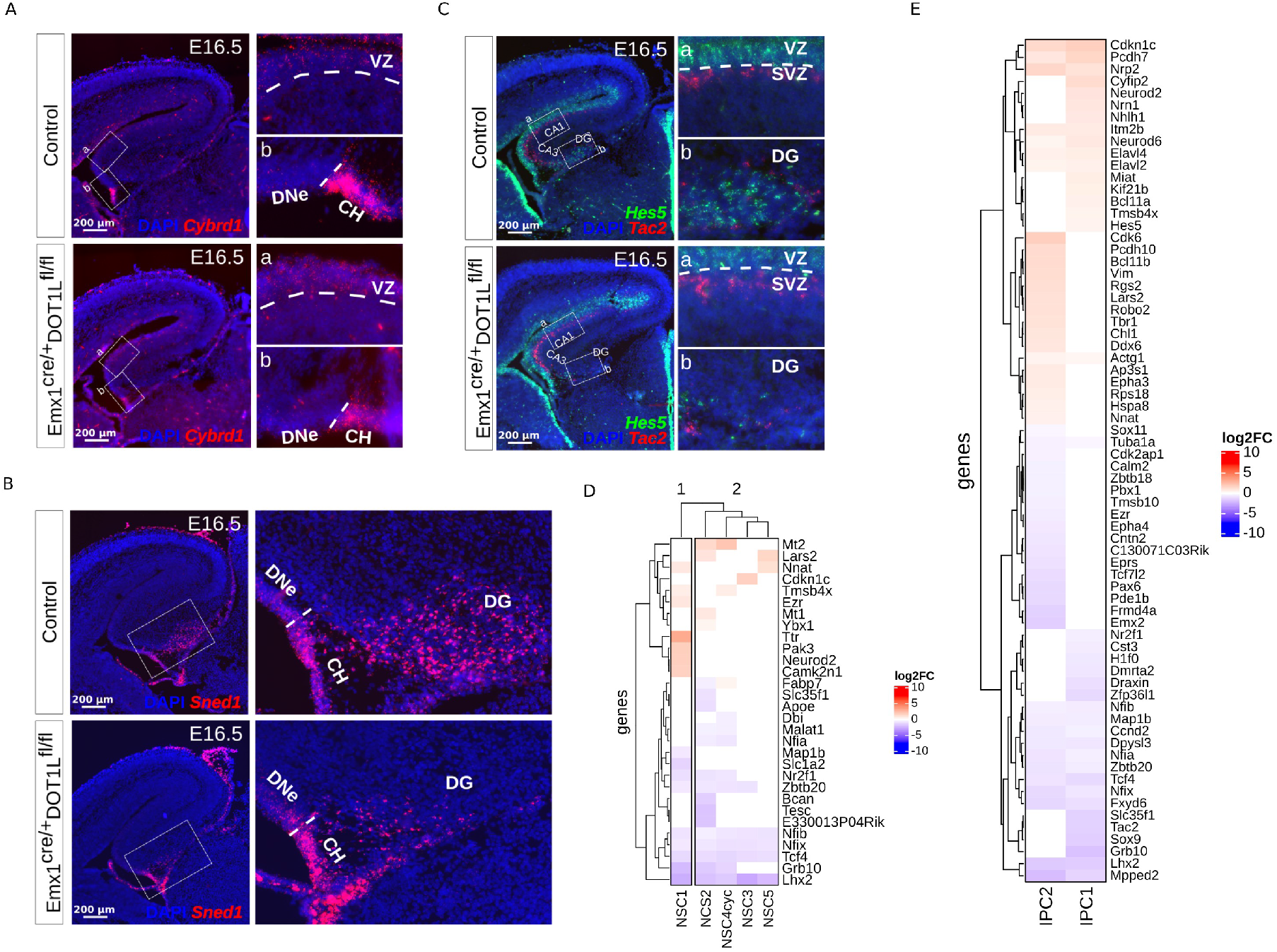
DOT1L is required for granule cells lineage generation and for controlling progression of IPCs towards differentiation. A-C) smFISH stainings for selected markers showing that NSC from the CANe are preserved (A) while granule cell lineage-related NSC (B) and IPC (C) are decreased upon DOT1L depletion (n = 3). Scale bars are indicated inside each image panel. DNe: dentate neuroepithelium; DG: dentate gyrus; VZ: ventricular zone; SVZ: subventricular zone; CH: cortical hem. D-E) Heatmaps of the DEGs (adjusted P<0.05) between conditions for the NSC clusters (D) and IPC clusters (E). Scale corresponds to log2 fold change between average expression for each condition within a cluster. Blue and red colors indicate significantly decreased or increased expression between DOT1L-cKO and control, respectively. Genes no significantly DE are depicted in white color.

NSC1 and IPC1 in the DNe/DG region seemed to be particular sensitive to DOT1L presence. We thus explored expression changes upon DOT1L-cKO in stem cells, which revealed that NSC1 separated from the other populations (Figure 4D). Seven genes altered significantly their expression level specifically in NSC1 but not in other stem cells (adjusted P<0.05). The increased expression of *Pak3*^38^ and *Neurod2*^39–41^ indicated that NSC1 were prone towards premature differentiation upon DOT1L-cKO. Among the IPCs, IPC1 had 19 unique DEG upon DOT1L-cKO compared to controls. Here, the decreased expression of *Nr2f1*^42^ and *Dmrta2/Dmrt5*^43^ correlated with precocious neuronal differentiation, similar to increased expression of *Cyfip2*^44^, *Neurod2*^39–41^ and *Nrn1*^45^ (Figure 4E). Thus, expression changes upon DOT1L-cKO suggested that premature differentiation caused the impaired development of the DG region.

### DOT1L prevents maturation and activates TFs involved in hippocampus development

We analysed DEGs within the entire data set to retrieve mechanistic insight into the observed phenotypic changes. 21 clusters contained cells from both experimental conditions and had DEGs (adjusted P<0.05), of which DG GC, CA1 and CA3 PCs had the highest numbers of DEGs (Figure 5A). Hierarchical clustering based on the transcriptional alterations upon DOT1L-cKO divided the cell populations into two main groups. Mature hippocampus cells (IPC2, DG GC, CA1 and CA3 PC) had evident changes in gene expression (Figure 5B**, group 2**). GO-term analysis of genes with decreased expression enriched in processes linked to brain development and morphogenesis. Genes with increased expression upon DOT1L-LOF retrieved significant enrichment for biological processes indicative of differentiation, including neuronal differentiation, synaptic and cognitive function terms (*q-value* < 0.05) (Figure 5C). From this finding we concluded that DOT1L impacted hippocampus development beyond the stem cells.

**Figure 5.**
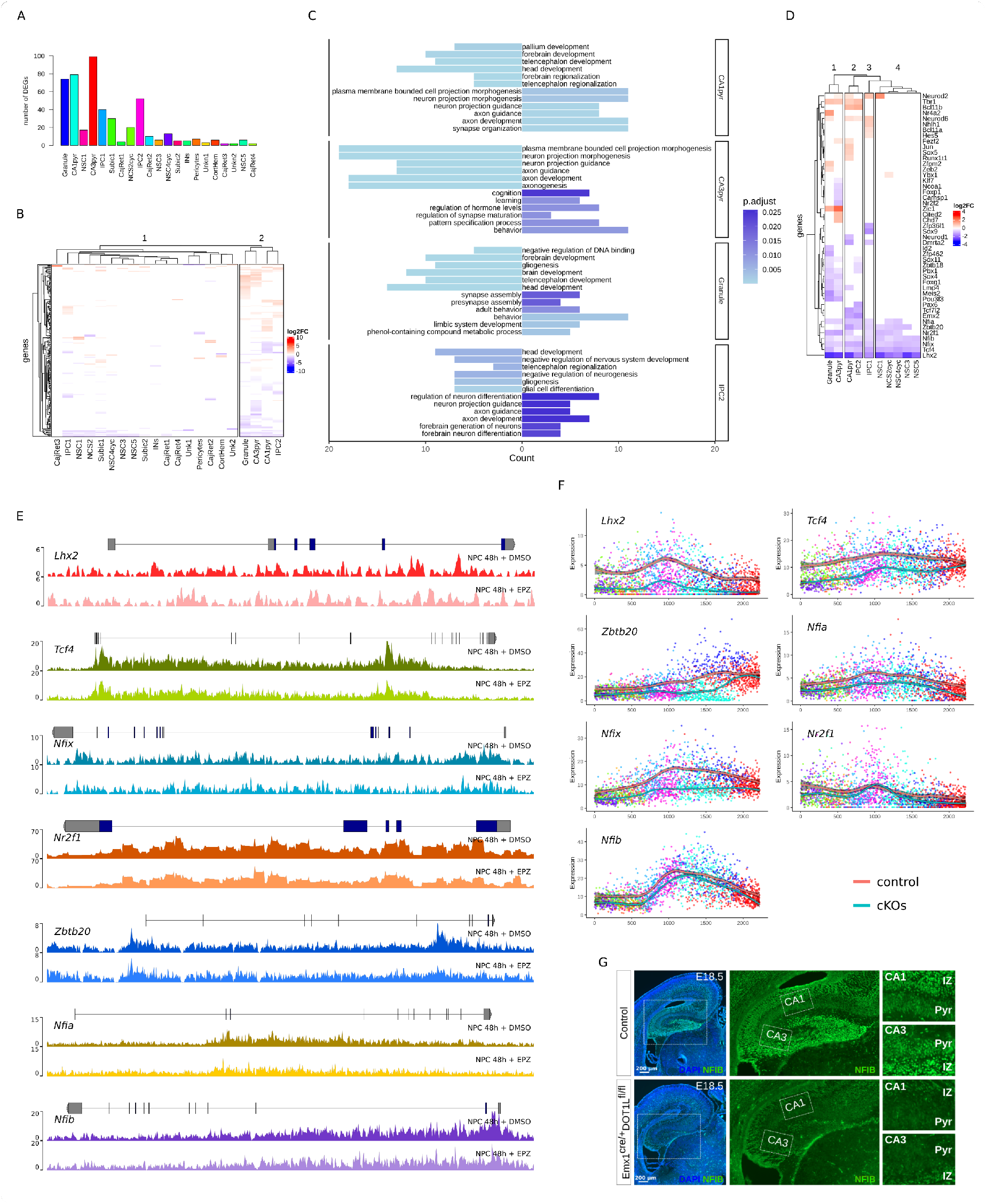
DOT1L prevents differentiation and activates master TFs involved in hippocampal development. A) Barplot showing the number of DEGs (adjusted P<0.05) obtained for the different cell clusters upon DOT1L deletion. B) Heatmap of the DEGs (adjusted P<0.05) between conditions for all the compared clusters. Scale corresponds to log2 fold change between average expression for each condition within a cluster. Blue and red colors indicate significantly decreased or increased expression between DOT1L-cKO and control, respectively. Genes no significantly DE are depicted in white color. C) Barplots of the enriched biological processes (*q-value* < 0.05) obtained for the gene sets significantly increased or decreased in expression in the respective clusters upon DOT1L-cKO. D) Heatmap of the transcription factors significantly increasing or decreasing in expression (adjusted P<0.05) in the DOT1L-cKOs for the clusters corresponding to the hippocampal lineage. Scale corresponds to log2 fold change between average expression for each condition within a cluster. Colour scale as in B. E) H3K79me2 profiles for *Lhx2, Tcf4, Nfix, Nr2f1, Zbtb20, Nfia* and *Nfib* in NPCs treated with DMSO (control) or EPZ for 48h showing decreased methylation signal in the gene bodies after EPZ treatment. Gene structures are shown on the top of the first track for each gene. F) Expression profiles for the genes in E along pseudotime axis. Red and light blue lines represent the smoothed running mean expression for control and cKO cells, respectively. Grey shadow along each line indicates a confidence interval of 0.95. Clusters are color coded as in Figure 1C. G) Immunofluorescence staining for NFIB on E18.5 control and mutant brain sections (n = 3) showing depletion of positive cells in the cKOs in different areas of the hippocampal region. CA: cornu ammonis; Pyr: pyramidal cell layer; IZ: intermediate zone. Scale bars are shown inside the image panels.

We focussed further analyses on TFs, because they have essential roles in hippocampus development. Expression levels of 49 TFs changed significantly upon DOT1L-cKO (adjusted P<0.05) (Figure 5D) and NSCs shared a decreased expression of known key regulators (Figure 5D**, cluster 4**). TFs both de- and increased in expression in mature cell populations. Similar TF expression grouped DG GC and CA3 PCs in cluster 1, and IPC2 and CA1 PCs in cluster 2. Noticeable, *Lhx2, Tcf4, Nfix*, and *Nfib* decreased in all, and *Nr2f1, Zbtb20* and *Nfia* in most represented populations. The genomic regions coding for these different TFs showed decreased H3K79me2 in neuronal progenitor cells (NPCs) treated with the DOT1L inhibitor EPZ5676, compared to DMSO control (Figure 5E), rendering this set of TFs as potential targets of DOT1L. All TFs also showed lower expression levels in the DOT1L-cKO along the pseudotime axis when compared to controls. *Lhx2, Nfix* and *Nfib* increased in expression from NSCs to IPCs, and decreased as the cells differentiated (Figure 5F). These results suggested a main role of the mentioned TFs in IPCs, in which chromatin remodelling gained particular relevance (Figure 2E, Node 9). A decreased expression on the protein levels of ZBTB20, LHX2 and NFIB (Figures 1F,1H, 5G) confirmed that DOT1L regulated these TFs involved in hippocampus development. Deletion of all the individual TFs impaired hippocampus development in a similar way as we observed^12, 19, 46^. This suggested a strong mechanistic link between altered TFs expression and the observed phenotype upon DOT1L-cKO.

### CA PCs are in two maturation states at E16.5

As CA PCs maturation lacks detailed understanding, and our data indicated that DOT1L-LOF impacted both cell populations transcriptionally, we characterized the differentiation trajectories of CA1 and CA3 PCs at single-cell resolution. We used the expression of *Pou3f1* and *Grik4*^6^, to annotate both PC clusters. Visual inspection of the data showed that neither *Pou3f1* nor *Grik4* was expressed homogeneously in all cells of the respective cluster. In contrast, CA1 and CA3 PCs contained cells with either high or low expression of *Pou3f1* or *Grik4*, which was indicative of unresolved heterogeneity within these two cell populations (Figures 6A,G). We performed independent sub-clustering analysis in each CA population (see methods) and resolved different maturation states. For both CA populations, one of the obtained sub-clusters enriched with cells expressing noticeable levels of either *Pou3f1* or *Grik4* (Figures 6B,H). GO-term analysis for significantly increased genes (adjusted P<0.05) in the CA1 and CA3 sub-clusters, enriched in either *Pou3f1* or *Grik4* expression, retrieved terms indicative for neuronal differentiation (Figures 6C,I). scvelo analysis corroborated that sub-cluster 1 cells (low *Pou3f1* (CA1) or *Grik4 (*CA3)) moved towards the sub-cluster 2 (high *Pou3f1* or *Grik4*), and supported the existence of two maturation states in CA1 and CA3 PCs at E16.5 (Figures 6D,J).

**Figure 6.**
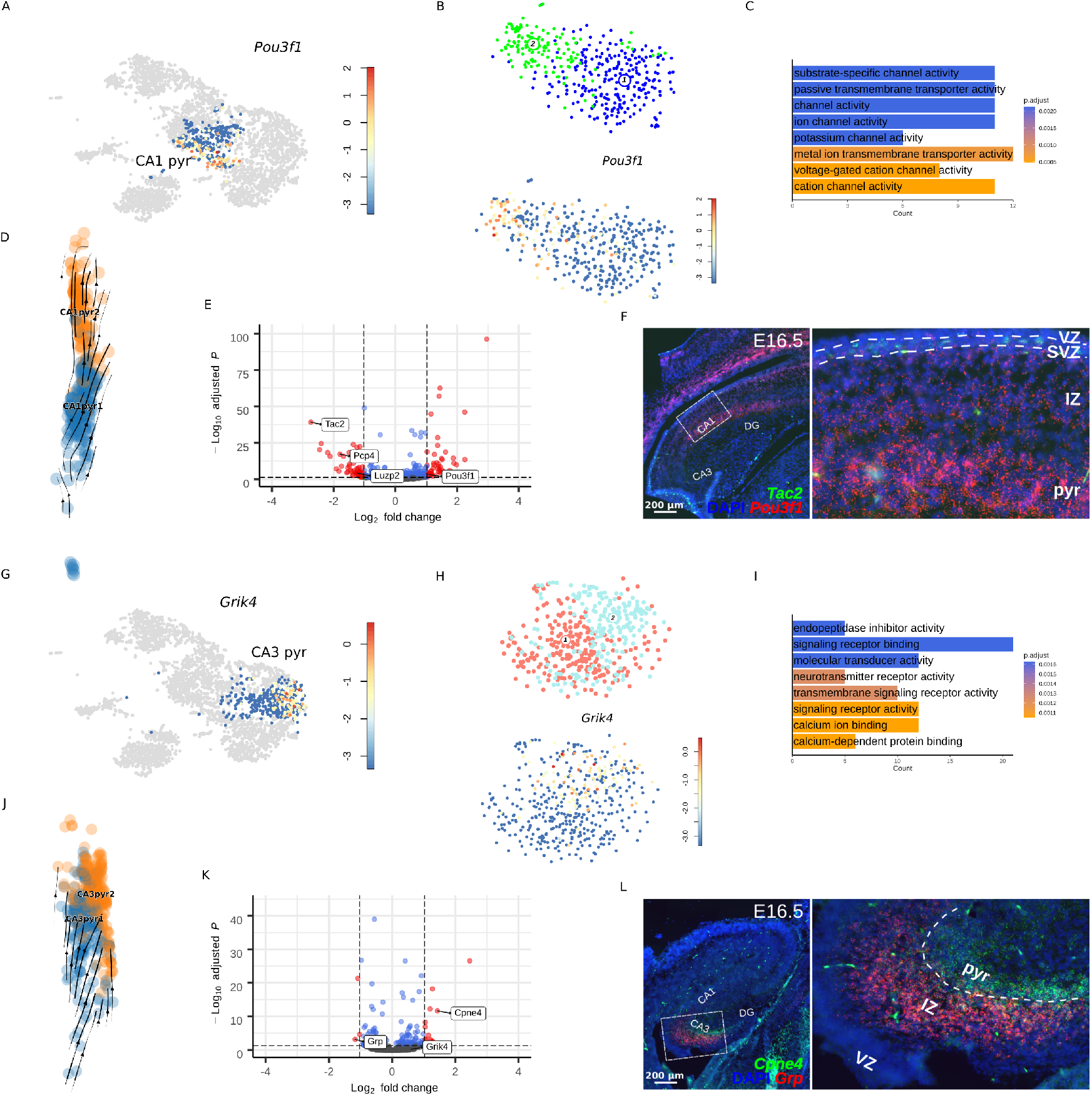
CA pyramidal cells are in two distinguishable maturation states at E16.5. A) and G) tSNE heatmaps showing non uniform expression of *Pou3f1* in CA1 pyramidal cells (A) and *Grik4* in CA1 pyramidal cells (G) (red: high; blue: low). Scale bars represent log2 values. B) and H) tSNE plots showing the 2 sub-clusters obtained after subclustering analysis (top) and apparent enrichment of *Pou3f1* (B, bottom) or *Grik4* (H, bottom) expression in the sub-cluster 2. Scale bars represent log2 values. C) and I) Barplots of the enriched molecular functions (*q-value* < 0.05) obtained for the gene sets significantly increased in expression in sub-cluster 2 for each CA region. D) and G) scvelo stream plots showing averaged velocity vectors (arrows) indicative of CA sub-cluster 1 cells movement towards sub-cluster 2. E) and K) Volcano plots for the DEGs (adjusted P<0.05) between sub-cluster 2 and sub-cluster 1 highlighting putative markers for the more mature (right) and less mature (left) cell states. F) and L) smFISH on E16.5 control brain sections (n = 3) for selected cell state marker genes. Scale bars are shown inside the image panels. CA: cornu ammonis; Pyr: pyramidal cell layer; SVZ: subventricular zone; VZ: ventricular zone; IZ: intermediate zone.

Based on different expression levels in the respective sub-clusters, we selected *Tac2, Luzp2* and *Pcp4* as markers for less mature and *Pou3f1* for mature CA1 PCs; *Grp* for immature and *Cpne4* for mature CA3 PCs, respectively (Figures 6E,K). Immature CA1 markers were expressed in the CANe, within the NSC3, 4cyc, and 5 expression domain, but their expression extended also into cortical areas including the subiculum. *Tac2* and *Pcp4* expression comprised the DNe/DG (Figures 6F**, S5**). *Pou3f1* expression extended in the IZ, where its expression was lower than in the overlying PCs, where mature cells expressed strongly *Pou3f1* (Figure 6F). *Grp* (immature CA3) was expressed in the CA3 region with a slightly higher level in the IZ than in PCs (Figure 6L). *Cpne4*, marking mature CA3 cells, was restricted to the PCs (Figure 6L). Together, we resolved two different maturation states during embryonic CA-field development that sequentially follow each other along the radial axis.

### DOT1L gates CA pyramidal cell maturation

DOT1L gates differentiation of progenitor cells, seemingly in a cell-type specific manner, e.g. with the CANe being less affected than the DNe. The transcriptional changes upon DOT1L-cKO in the mature CA1 and CA3 PCs (Figures 5A-C) and the two different maturation states in both CA-fields, suggested that DOT1L gates also maturation of differentiated cells. Indeed, DOT1L-LOF favoured the transition from less to more mature PC states, because more cells belonged to mature CA1 or CA3 sub-clusters compared to controls, which contained a significantly higher proportion of cells in less mature sub-clusters (adjusted P<0.05, Fisher’s test) (Figures 7A,G). In support and indicative of premature differentiation, mutants showed a stronger *Pou3f1* signal at the level of the SVZ and IZ compared to controls (Figure 7B). The CA3 marker *Grp* was nearly completely lost in DOT1L-cKO, corroborating a drastic depletion of the less mature cells (Figure 7H). DEG analysis revealed discrete gene sets, expression of which either decreased or increased significantly in the less mature cell states upon DOT1L-cKO compared to controls (adjusted P<0.05) (Figures 7C,I). In both CA-fields significantly decreased gene sets enriched for GO-terms related to transcriptional control (*q-value*<0.05), which indicated that TFs represented again important targets affected upon DOT1L-cKO (Figures 7C,I). We determined, independent on the DOT1L-cKO, TFs that changed significantly in expression in mature compared to less mature CA cells (Figures 7D,J). Both, CA1 and CA3 maturation correlated with decreased expression of *Lhx2*, *Nfia, Nfib, Nfix*, and *Nr2f1* (adjusted P<0.05) (Figures 7D,J). The significant higher expression of the former TFs in the less mature cell state and their role as transcriptional activators indicated that they are favouring a less differentiative cell state. We subsequently identified those TFs that might mediate the increased maturation upon DOT1L-cKO by intersecting the TFs changing upon CA maturation with TFs changing significantly upon DOT1L-cKO in less mature CA cells. 13 TFs for CA1, and 8 for CA3 were differentially expressed (adjusted P<0.05) (Figures 7E,K), of which *Nfia, Nfix, Nr2f1* and *Lhx2* decreased, and *Bcl11b* increased in expression upon DOT1L-LOF in both CA-fields (Figures 7F,L)*. Lhx2* was transcriptionally most affected and its expression also decreased on the protein level in the CANe, IZ and the pyramidal layers upon DOT1L-cKO (Figures 7M, 1H).

**Figure 7.**
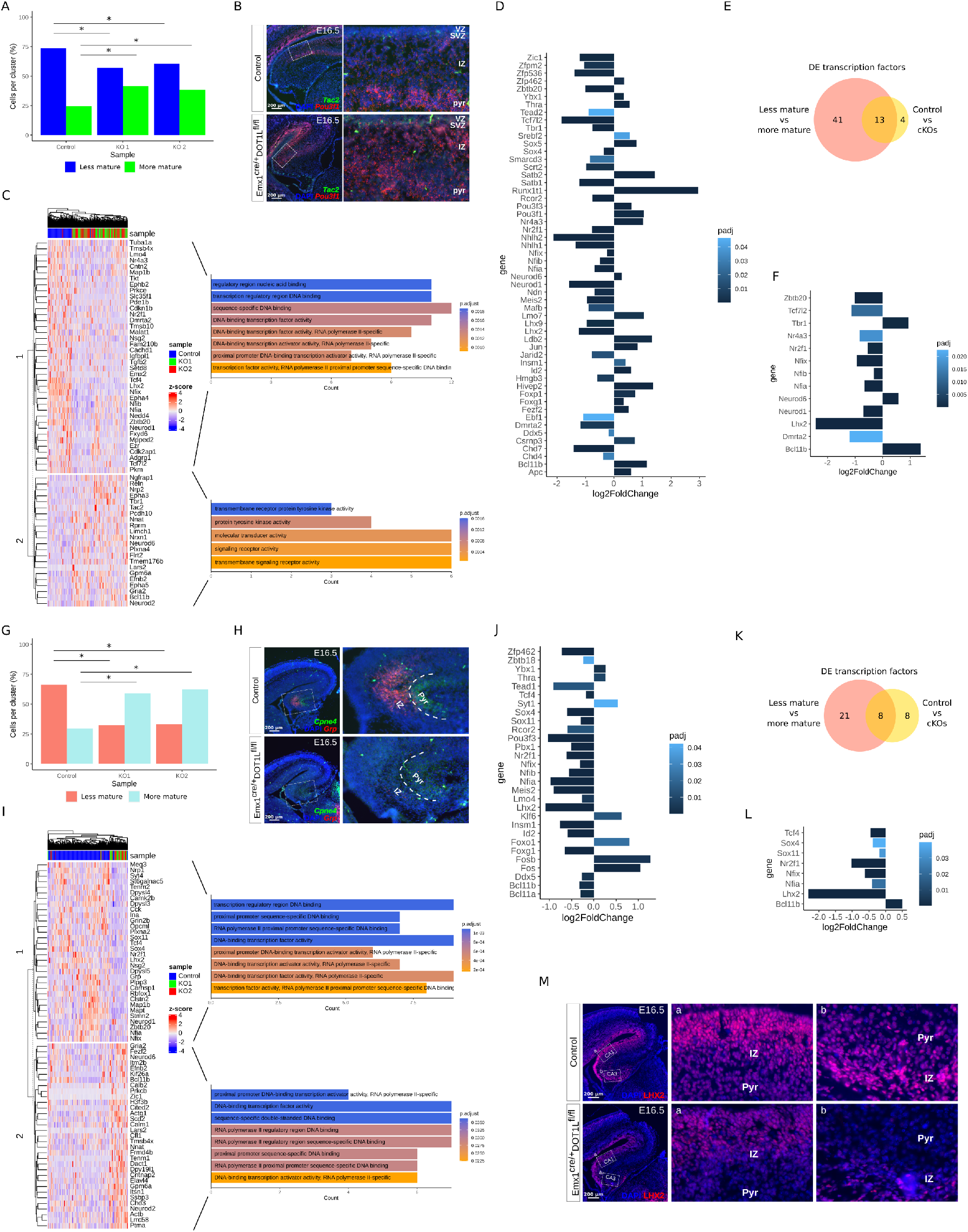
Loss of DOT1L promotes maturation of CA pyramidal cells. A) and G) Barplots of cell proportions in each sub-cluster, normalized by sample. Significant changes in cell proportions based on Fisher’s exact test (adjusted P<0.05) are indicated with asterisks. Colour codes are kept as in Figure 4 B and H (top). B) and H) smFISH on E16.5 control and cKO brain sections (n = 3) for the selected cell state gene markers. C) and I) Heatmaps (left) showing the DEGs (adjusted P<0.05) between control and cKO samples in the less mature CA sub-clusters. Genes are grouped based on hierarchical clustering and divided in 2 main clusters according to the direction of change. Barplots (right) indicating the main biological processes enriched (*q-value*<0.05) in each of the 2 gene clusters are shown. D) and J) Barplots for the differentially expressed TF (adjusted P<0.05) between the 2 CA maturation states. Genes changing in expression in the more mature state relative to the less mature state are shown. E) and K) Venn diagrams indicating the intersection of the TFs differentially expressed between more and less mature sub-clusters with TFs differentially expressed between conditions for the less mature cell state. F) and L) Barplots showing the log2 fold change in expression for the differentially expressed TF (adjusted P<0.05) in the less mature cell state upon DOT1L depletion that are also differentially expressed between maturation states. M) Immunofluorescence staining for LHX2 on E16.5 control and mutant brain sections (n=3) showing depletion of positive cells in the IZ and Pyr upon DOT1L-cKO. Scale bars are shown inside each image panel.CA: cornu ammonis; Pyr: pyramidal cell layer; VZ: ventricular zone; SVZ: subventricular zone; IZ: intermediate zone.

## Discussion

The presented data provide novel insights into cell lineage trajectories and fates in the developing hippocampus that are under control of the histone methyltransferase DOT1L. DOT1L functions in the CNS to gate and control adaption of subsequent cell fates^22–24, 47^ and LOF mice had an impaired hippocampal structure^22^. We used the DOT1L-cKO to dissect cell lineage differentiation in the hippocampus that is in this regard remarkably understudied. Based on high-resolution scRNAseq, we demonstrated different stem cell populations in the developing hippocampus. Other recent work that applied scRNAseq technology for the study of the developing hippocampus represented NSCs as one homogeneous population, and did not discriminate either between NSCs and IPCs committed to specific hippocampal lineages (i.e., PCs and GCs) or between possible distinct NSC states^5, 32^. We resolved heterogeneity in the stem cell populations of the hippocampus. The subdivision of NSCs and IPCs into discrete clusters pointed either towards transitory cell states during proliferation and differentiation, or to discrete progenitor classes with the potential to generate specific cell lineages. These alternatives were not exclusive, but were both true. We defined a set of novel marker genes to discriminate the two germinative regions, CANe and DNe, that were proposed by others, but were molecularly ill-defined due to lack of specific markers for the respective cell populations. From recently reported markers, used to dissect DG development on a protein level (SOX9, HES1/5, NOTCH1/2, HEY2 for NSCs; EOMES, DLL3 for IPCs^10^), only *Dll3* enriched in one single specific cell population in our data set, i.e. IPC1. All other markers were transcribed at similar levels in all identified NSC clusters in our study (**table 1**), and, moreover, some of them in Cajal-Retzius cells, oligodendrocyte precursors, CH or pericytes. *Eomes* was expressed by NSC1 and 3, by Cajal-Retzius cells and in both IPC clusters. Thus, our data is of major relevance for high resolution studies of lineage relationships in the developing hippocampus. We identified NSC3, NSC4cyc, and NSC5 as CANe, expressing *Sox21* and *Cybrd1*, the NSC2cyc as DNe, expressing *Sned1* and *Adamts19*, and NSC1 feeding the DG trajectory. Given that derailed development of the hippocampus is underlying diseases like autism and schizophrenia^48, 49^, it is of paramount relevance to deepen our knowledge on the basic cellular relationships in this part of the limbic system.

Our data advance prevailing views on hippocampus development. Firstly, our data analysis suggested that subiculum cells lacked connection to NSCs and IPCs present in the E16.5 hippocampus. This points towards an extra-hippocampal origin of subiculum cells, which argues against recent findings that subiculum and CA1 PCs share a common developmental path^32^. The discrepancy between both studies could be attributed to the different developmental stages analyzed (E16.5 in our work, P0 and P5 in La Manno et al^32^). Moreover, the bioinformatic tools (scvelo vs Velocyto) and their specific parameterizations (for example selection of variable genes) employed to predict lineage relationships might also affect the predictions^50^.

Secondly, we observed that the CA1-lineage was affected less severely compared to CA3 upon DOT1L-cKO. A possible reason for this could be that both populations originate from, however so far not defined, different progenitor pools of which only one is DOT1L-sensitive. Our data set could suggest a trajectory from NSC1 via IPC1 to CA3 and DG, because these clusters decreased in cell numbers upon DOT1L-cKO. A common CA3/DG progenitor could be postulated as different concentrations of WNT3A differentiated human induced pluripotent stem cell-derived hippocampal progenitors into either CA3 PC or DG GC^51^. Moreover, *Prox1-*LOF in postmitotic neurons induced transcriptional changes indicative of a switch from DG GC to CA3 PC identity^52^, highlighting an inter-conversion plasticity between both lineages. Despite the quantitative changes observed in our data set and the transcriptional similarity among CA3 PCs and DG GCs, the algorithms we used to predict this lineage trajectory in the scRNAseq data set and our visualisations in tissue sections could not find support for CA3 PCs deriving from the DNe. Because, surprisingly, our in-depth analysis of differentiation trajectories into CA1 and CA3 PCs, DG GCs or subiculum cell fates did not reveal convincing differentiation probability from any NSC/IPC clusters towards CA3. The lack of identification of CA3 progenitors could be caused by technical limitations, but our data set derived from three individuals. The likelihood that we did not capture CA3 progenitors in three different cell isolation attempts seems low. This raises the hypothesis that CA3 progenitors are not represented at E16.5, implying that they occur earlier in development. This is possible, because CA-field development is controlled temporally, with CA3 arising earlier compared to CA1^8^. Our data supports this observation, as at E16.5 CA3 PCs were the most differentiated cell population. Thus, CA3 progenitors should differentiate earlier than CA1 progenitors, if they share a common ancestor. Or, equally possible, CA3 progenitors differentiate earlier but from a shared ancestor of the DG granule lineage. Precise definition of lineage trajectories and spatio-temporally separated progenitor populations for the CA1- and CA3-fields requires time resolved scRNAseq experiments. Using DOT1L-cKOs with relative normal numbers of CA1 but loss of CA3, seems an adequate approach to provide such in-depth insights.

Thirdly, previous studies proposed maturation of the CA PCs along the longitudinal axis in a pole-inward pattern, starting at embryonic and prolonging into postnatal stages^6^. We refine this view by showing two radially separated maturation states in the CA PC lineages. *Grp* and *Cpne4* are highly specific novel markers that highlight this radial maturation trajectory of CA3 PCs. In conjunction are altered expression levels of TFs, most of which decreased with maturation. DOT1L-LOF decreased a TF-set that is silenced during normal developmental CA-field maturation, which might be responsible for the advanced maturation observed in both CA-fields. Additionally, a set of TFs is differently regulated during the maturation of CA1 or CA3 PCs: *Zfp462, Pou3f3, Insm1, Id2, Foxg1* and *Bcl11b* increased during CA1-maturation, and they decreased during CA3-maturation (Figure 6D,J). Thus, this TF-set might be involved in cell-specific maturation programs, and it might confer the loss of CA3 and relative normal numbers of CA1 PCs upon DOT1L-LOF. In this regard DOT1L-cKO increased *Bcl11b* expression in CA3 PCs, thus in opposing direction compared to the normal maturation trajectory. *Bcl11* homologues might play important roles in conferring lineage identity between CA-fields because whereas *Bcl11a* and *Bcl11b* are co-expressed in CA1/2, *Bcl11b* is not expressed in CA3, but *Bcl11a* expression extends into this region^53^. Increased levels of *Bcl11b* upon DOT1L-cKO specifically in the CA3 PCs might interfere at least in part with adapting of a phenotypically distinct CA3-identity.

Lastly, our data establish DOT1L as first histone methyltransferase to act as an upstream regulator of instructive TFs that maintain progenitor states, but also drive maturation of CA PCs or DG GCs. The histone deacetylases, HDAC1, 2 and 3^54–56^ are other histone modifiers impacting hippocampus development. Thus, epigenetic mechanisms play pivotal roles in hippocampus development, in cell fate and maturation processes. Our data suggest that chromatin plasticity and histone modifications are important mechanisms for generating and resolving transient developmental stages in the hippocampus.

## Methods

### Generation of DOT1L-cKO mice, dissection and genotyping

All animal experiments were approved by the animal welfare committees of the University of Freiburg and local authorities (G16/11). Emx1-cre^57^ and Foxg1-cre^58^ mice were kept on a C57BL6/J background and used to breed Emx1-Dot1l and Foxg1-Dot1l-cKO mice, respectively. Mutant offspring correspond to both Emx1^cre/+^;Dot1l^fl/fl^ and Foxg1^cre/+^;Dot1l^fl/fl^, while WT offspring correspond to Emx1^+/+^;Dot1l^fl/fl^, Emx1^+/+^;Dot1l^fl/+^, Foxg1^+/+^;Dot1l^fl/fl^ and Foxg1^+/+^;Dot1l^fl/+^.

Adult pregnant animals were sacrificed by cervical dislocation for embryo collection at embryonic day (E) E16.5 or E18.5. The embryos were sacrificed by decapitation. The brains isolated at defined embryonic stages were washed in cold phosphate-buffered saline (PBS) and then fixed in 4% PFA overnight at 4° C. For cryosections, fixed brains were incubated in 30% sucrose solution at 4° C until, embedded in tissue freezing medium (Leica Biosystems, Germany) and frozen at −80 °C. Forebrains were cut coronally into 16μm sections and mounted on Superfrost Plus Microscope Slides (Thermo Fisher, USA). Genotyping was carried out by polymerase chain reaction (PCR) from tail DNA. Briefly, tail samples were lysed in QuickExtract DNA Extra Solution 2.0 (Lucigen, USA) and PCR reactions were carried out with GoTaq DNA polymerase and with the Primers: Emx1-Cre forward: 5’-ATGCTTCTGTCCGTTTGCCG-3’, Emx1-Cre reverse: 5’-CCTGTTTTGCACGTTCACCG-3’, DOT1L forward: 5’-GCCTACAGCCTTCATCATTC-3’, DOT1L reverse: 5’-CCCATACAGTACTCACCGGAT-3’, Bf1-F25 forward: 5’-GCCGCCCCCCGACGCCTGGGTGATG-3’, Bf1-R159 reverse: 5’-TGGTGGTGGTGATGATGATGGTGATGCTGG-3’ and Bf1-Rcre222 reverse: 5’-ATAATCGCGAACATCTTCAGGTTCTGCGGG-3’. PCR products were analyzed by agarose gel electrophoresis.

### Histological analyses

#### Cresyl Violet (Nissl) Staining

Frozen embryonic brains were sectioned at 16µm with a Leica cryostat (CM3050S) and stained with cresyl violet, following standard protocol. The sections were mounted on Superfrost Plus Microscope Slides (Thermo Fisher, USA) with Eukitt mounting medium (O. Kindler, Germany).

#### Immunofluorescence staining (IF)

Embryonic mouse forebrain sections fixed on Superfrost Plus Microscope Slides (Thermo Fisher, USA) were surrounded with a hydrophobic pen and permeabilized and blocked with 10% normal donkey serum (NDS; Bio-Rad, USA)/ 0.1% Triton X-100 (Carl Roth, Germany)/ PBS for 1 h at room temperature (RT). Sections were incubated with primary antibodies diluted in the blocking solution over night at 4° C. After 3 washing steps with PBS/ 0.1% Triton X-100 the secondary antibodies (dilution 1:500) were applied in blocking solution and incubated for 1 h at room temperature. Sections were washed again 3 times with PBS and nuclei were stained with 4’,6-diamidino-2-phenylindole (DAPI, dilution 1:1000). After 3 final washing steps sections were mounted with Dako fluorescent mounting medium (Agilent Technologies, USA) and sealed with nail polish. The respective antibodies and the dilutions used are listed in **table 2**.

#### In situ hybridization (ISH)

Forebrain slices at the different developmental stages were hybridized with digoxigenin-labelled riboprobes in hybridization buffer (12.7 mM Tris base, 184.4 mM NaCl, 5.9 mM NaH_2_ PO_4_, 6.27 mM Na_2_ HPO_4_, 5 mM EDTA pH 8.0, 0.5x Denhardt’s solution, 1 mg/ml Yeast RNA, 10% Dextran sulfate, 50% v/v Formamide) at 68°C overnight. Sections were washed 3 times in a solution containing 50% formamide, 0.1% Tween-20 and 5% saline sodium citrate at 68°C in a water bath. They were then transferred to an incubation chamber and washed twice with maleic acid buffer and Tween-20 (MABT) for 30 min at RT. After blocking in a MABT solution containing 20% lamb serum, sections were incubated with an alkaline phosphatase-conjugated anti-digoxigenin antibody (1:1500 in blocking solution; Roche, Switzerland) overnight at RT. After four washing steps in MABT (10 min, 3x 20 min) and three washing steps (7 min each) in pre-staining buffer, the reaction product was developed using NBT/BCIP solution diluted in pre-staining buffer (1:100; Roche, Switzerland) overnight at RT. Stained sections were washed 4 times in PBS and then embedded using Aquatex (Merck Millipore, USA).

#### Single molecule FISH (smFISH)

Brain tissue cryosections with a thickness of 16µm were processed following the RNAScope kit protocol, with some few modifications. Firstly, the tissue sections were incubated at 40°C for 1 h in the HybEZ hybridization oven (Advanced Cell Diagnostics, USA) to ensure adhesion of the tissue on the slides. The sections were then washed 5 min in 1X PBS to remove the remaining tissue freezing medium. Afterwards tissue slides were treated with RNAscope Hydrogen Peroxide for 10 min at RT and the sections were immediately washed in deionized water. Secondly, the slides were put in a slide holder and incubated for 5 min inside a beaker filled with 1X RNAscope Target Retrieval Reagent preheated at approximately 95°C. At the end of the incubation time the slides were washed in deionized water, incubated in 100% Ethanol for 2 min, and then dried at RT. Afterwards, an hydrophobic pen was used to delineate the area around the sections and RNAscope ProteaseIII was added on top of the sections followed by incubation 30 min at 40°C in the HybEZ hybridization oven. Thirdly, the slides were washed with deionized water and subjected to the corresponding hybridization and amplification steps in the HybEZ hybridization oven at 40°C. At this step the sections were incubated with the specific probes for 2 h. For the amplification, the sections were incubated with RNAscope Multiplexv2 Amp1 for 30 min, RNAscope Multiplexv2 Amp2 for 30 min and RNAscope Multiplexv2 Amp3 for 15 min, each step followed by two 2 min washing step on Washing Buffer at RT. Fluorescent signal was revealed by incubation of the sections with RNAscope Multiplex FLv2 HRP-C specific reagents (C1, C2 or C3 depending on the probes channel) for 15 min, the respective fluorophore for 30 min and then RNAScope Multiplex FL v2 HRP Blocker for 15 minutes, each step followed by two 2 min washing step on Washing Buffer at RT. In the case of double stainings the slides were incubated with the respective RNAscope Multiplex FL v2 HRP-C reagent targeting the second probe and the respective fluorophore following the same steps as above. All the incubation steps at 40°C were carried out inside an aluminium chamber containing the slides arranged on an ACD EZ-Batch Slide Rack and a deionized water moistened tissue. Finally, the slides were counterstained with DAPI for 30 sec at RT and mounted using Dako fluorescent mounting medium (Agilent Technologies, USA). The dyes used for signal detection were Opal Dye 520 and Opal Dye 570 (Akoya Biosciences, USA). The respective probes as well as the fluorophore dilutions used in the different stainings are listed in **table 3**.

#### Image acquisition and processing

Images were acquired with Axio Imager M2 (Zeiss, Germany). High resolution images were downscaled using GIMP (version 2.10.24) and figure panels were arranged using Inkscape (version 1.0.2: 394de47547, 2021-03-26).

### scRNAseq experiment

#### Sample size estimation

To determine the number of cells that we needed to sequence for stable clustering, we used data from the cortex, hippocampus and subventricular zone of two E18 mice (https://support.10xgenomics.com/single-cell-gene-expression/datasets/1.3.0/1M_neurons). The data set was filtered for hippocampal cell populations based on the expression of marker genes, namely *Zbtb20, Satb2, Etv1, Man1a, Pou3f1, Prox1, Pax6 and Mef2c*. The obtained data set served as a pilot data set to determine the number of cells for the planned experiment. We used cells from the original (filtered) dataset to determine cluster stability with varying numbers of cells (250, 500, 1000, 2000, and 3000). To determine the clustering stability for a higher number of cells, exceeding the number of cells in the pilot data, we trained the single-cell deep Boltzmann machine (scDBM)^26^ on the pilot data set and generated synthetic cells which we combined with the initial data.

The number of cells at which the clustering reached stability, meaning that there were no or few outlier clustering solutions, was chosen as the optimal number of cells.

First, we subsampled 250, 500, 1000, 2000, and 3000 cells from the pilot data for 30 times each and ran k-means clustering on each of these subsamples. From previous analyses, we saw that 16 clusters were appropriate for the pilot data set, and therefore we defined k = 16 for the clustering. Next, we assessed clustering stability for each subsample using the Davies-Bouldin Index (DBI). Additionally, we trained a scDBM on the pilot data. We generated 1000 and 2000 (30 times each) additional cells, after which we combined the synthetic cells with the pilot data and performed k-means clustering. A stable clustering occurred at a cell count of 4000 cells, indicated by a generally low DBI and few outliers after clustering the 4000 cells for 30 times.

#### Cell dissociation, sorting and sample processing for scRNAseq

Three samples corresponding to 1 control and 2 cKOs at stage E16.5 were collected and processed for scRNAseq analysis. The hippocampi of E16.5 embryos were dissected on cold HBSS (Hank’s Balance Salt Solution) buffer and transferred to 1.5 ml tubes containing the dissociation solution (Papain + DNase). The dissociation procedure was carried out with the Papain Dissociation System (Worthington-Biochem, USA) following the manufacturer’s instructions with some few modifications. Briefly, all the reagents were reconstituted in the respective solutions and equilibrated with 95% O_2_ : 5% CO_2_ if needed. The dissociation solution (Papain + DNase) was pre-incubated at 37°C and aliquoted in 1.5 ml tubes for sample collection. Tissue samples were disrupted inside the tubes using a p200 pipette tip precoated with normal goat serum and then incubated at 37°C in a water bath for approximately 30 min with manual inversion every 5 min. After the incubation time samples were pipetted up and down, as mentioned above, for obtaining a single cells suspension. The tubes were left undisturbed for 60 sec and the supernatants were transferred to another tube and pelleted down by centrifugation at 300 rcf, 4°C, 5 min. Supernatants were discarded and the cells were resuspended with resuspension solution (DNase + Ovomucoid inhibitor) using p200 pipette tips precoated with serum, and immediately and carefully transferred on top of another set of tubes containing Ovomucoid inhibitor solution. The tubes were then centrifuged at 76 rcf, 4°C, 6 min, the supernatants were removed, and the cell pellets resuspended in 200μl HBSS buffer and kept on ice.

The cells were stained with Zombie Green™ Fixable Viability Kit (BioLegend, USA) in PBS for 15 min at RT and in the dark using a dilution of 1:500. After that the cells were washed with 1ml 1x MojoSort™ Buffer (BioLegend, USA), centrifuged at 300 rcf for 5 min, resuspended in 2 - 3 ml 1x MojoSort™ Buffer and passed through a 30 µm filter (Sysmex, CellTrics). Finally live cells were sorted on a MoFlo XDP Cell Sorter (Beckman Coulter, USA) into 384-well plates containing prepared lysis buffer and processed with the CEL-Seq2 modified protocol (Sagar et al., 2018). Briefly, the single cells were sorted in 384 well plates containing RT (reverse transcription) primers (anchored polyT primers having a 6 bp cell barcode, 6 bp unique molecular identifiers (UMIs), a part of 5’ Illumina adapter and a T7 promoter), dNTPs, Triton X-100 and Vapor-Lock (Qiagen). The MoFlo XDP Cell Sorter was calibrated for dispensing the single cells in the centre of each well prior to sorting and the machine was run using trigger pulse width to exclude doublets. The 384 well plates containing the sorted cells were centrifuged at maximum speed and stored at −80°C until library preparation as described in Sagar et al., 2018. Libraries were sequenced on a HiSeq3000 Illumina sequencing machine and the demultiplexing of the raw data was performed by running bcl2fastq (version 2.17.1.14.).

#### scRNAseq data analysis

##### Quantification of transcript abundance

BWA (version 0.6.2-r126) was used to align paired-end reads to the transcriptome using default parameters (H. Li & Durbin, 2010). The transcriptome was based on all gene models from the mouse ENCODE VM9 release from the UCSC genome browser, which contained 57207 isoforms with 57114 isoforms mapping to fully annotated chromosomes (1-19, X, Y, M). Gene isoforms were merged per gene to a single gene locus and gene loci were grouped to gene groups, if there was >75% overlap, resulting in 34111 gene groups. The right mate of the read pair was mapped to all gene groups and multimapping reads were excluded. The left mate carried the barcode information with 6bp corresponding to the UMI and the following 6bp to the cell barcode. The remaining sequence contained a poly(T) and adjacent gene information but was not used for quantification. Transcript counts were obtained by aggregating the number of UMI per gene locus and for every cell barcode^59^.

##### Filtering, normalization, dimensionality reduction and clustering

The output count matrices generated were combined using R (versions 3.6.3 and 3.5.1) and Rstudio (version 1.2.5001, build 93 (7b3fe265, 2019-09-18)) and the data filtering, normalization, dimensionality reduction and clustering analyses were performed with the RaceID package (version 0.1.3). As pre-processing steps, the genes with names starting with “mt”, “Gm” or “Rik” were filtered out from the expression matrix, and cells with less than 500 total counts were removed. For the filtering step we ran RaceID filterdata function with default parameters after adjusting *minnumber=1, CGenes= c(“Mki67”, “Pcna”)* and *Fgenes=c(“Kcnq1ot1”).* A total of 3701 cells were kept after filtering (control = 1287; cKO1= 1102; cKO2= 1312) and used for further analysis. For the computation of the distance between cells we used the default RaceID settings and for the clustering we set the clustexp function parameter *cln=16*, which represents the number of clusters at which the change in the log of within cluster dispersion became small. Additionally, outlier estimation was performed using the default parameters.

##### Sub-clustering analyses

For sub-clustering analyses the count matrix was subset using the information of the cell names belonging to the specific clusters. The RaceID pipeline was run using as input the subset data and following the steps described above, setting *cln=*2 and the *metric=* “*logpearson”* as the metric used for computing the distance between cells. The selection of 2 as the expected number of clusters was based on the observed gene expression patterns on the t-SNE representations. We proposed that the 2 expected clusters should represent: less and more mature cells. On the other hand, we defined log-Pearson as the distance parameter for the clustering step. This selection was based on the idea that a putative unresolved heterogeneity could be driven by lowly expressed transcripts that were not considered for the clustering step in our initial analysis.

##### Analysis of the effect of sex and genotype on clustering

To ensure that there were no profound differences between mice of different sex that could potentially confound our analyses, we visualized the biological variation in the data via a denoised principal components analysis (PCA)^60^ using the scran package (version 1.16.0). Here, a threshold value for the number of principal components is calculated based on the proportion of variance explained by the biological components in the data. We separated male and female mice using the expression of *Xist* gene (2 males: 1 control and 1 cKO; and 1 cKO female). By visual examination of the first five principal components, we could not detect any substantial differences between the sex of the mice and thus ruled out the sex as confounding factor. A similar analysis performed using the genotype (control versus cKO) as grouping factor, which showed the absence of major effects due to the genotype.

##### Differential gene expression analysis

Analyses of the differentially expressed genes were performed with RaceID built in function. Briefly, a background model for the expected transcript count variability is internally computed by RaceID algorithm, and used for inferring negative binomial distributions reflecting the gene expression variability within each group. A P-value for the observed difference in transcript counts between the two compared groups is calculated based on the inferred distributions, and subsequently adjusted for multiple testing using the Benjamini–Hochberg method. Expression of the *Xist* gene was removed from the DEGs for the distinct plots generated.

##### Comparative analysis of cell proportions

Comparison of cell population proportions between control and cKOs per cluster was carried out in a 1 versus 1 basis using the Fischer’s exact test, by means of the fisher.test function included in R. Additionally, in order to correct for multiple comparisons a post hoc adjustment of the obtained Pvalues was performed using the p.adjust function with Bonferroni as the correction method. We considered a change in proportions between conditions for a specific cluster to be significant only if both cKO samples showed an adjusted P*-*value less than 0.05 when individually compared to the control sample.

### RNA velocity analysis

#### Single cell expression data analysis

##### Reference and Quantification

In order to estimate levels of both spliced transcripts and introns for use in RNA velocity analysis, a reference for pseudo-alignment was constructed similar to the “alevin_sep_gtr” method described^50^.

Mouse genome sequence and transcript annotation were obtained from Ensemble release 98 (fasta/mus_musculus/dna/Mus_musculus.GRCm38.dna.primary_assembly.fa and gtf/mus_musculus/Mus_musculus.GRCm38.98.gtf files from http://ftp.ensembl.org/pub/release-98/). The sequences of transcripts and introns were extracted using R (https://r-project.org, version 3.6) and the eisaR package (version 0.8 available from https://github.com/fmicompbio/eisaR/tree/051496c6f90a7a4320821a8f3c0518fb4cf4a85a) using extractTxSeqs with type = “spliced” and extractIntronSeqs with type = “separate”, flanklength = 50, joinOverlappingIntrons = FALSE. A transcript-to-gene table was created which links transcripts to genes, and introns to distinct genes for simultaneous quantification (see below). The whole procedure was then performed as described in http://bioconductor.org/packages/release/bioc/vignettes/eisaR/inst/doc/rna-velocity.html. Extracted sequences were indexed with Salmon (version 1.1.0)^61^ with arguments -k 23 -- type puff --gencode and using the genome as a decoy. Reads were quantified using Salmon/Alevin (version 1.1.0)^62^ with parameters --celseq2 and the transcript-to-gene map created above.

##### Quality control and filtering

Technical quality of single cell experiments, cell barcode identification and quantification were assessed using the Bioconductor package alevinQC (version 1.2.0, https://doi.org/doi:10.18129/B9.bioc.alevinQC). Salmon/Alevin counts from all samples were imported into R using the tximeta package (version 1.4.3)^63^ and stored in a SingleCellExperiment container for downstream analysis (55421 genes, 5678 cells). The number of detected genes per cell and the fraction of counts in genes encoded on the mitochondrion (chrM) were calculated using the addPerCellQC function from the scater package (version 1.14.6)^64^, and cells with more than 1000 detected genes and less than 25% mitochondrial counts were retained (3972 cells). Cell size factors for normalization of raw counts were calculated using the scran package (version 1.14.6)^60^, by first clustering the cells using the quickCluster function with method=“igraph” and then computeSumFactors with min.mean=0.5, which implements the deconvolution strategy described^65^. Log-transformed normalized counts were then calculated using the logNormCounts function from the scater package.

#### Velocity estimation

RNA velocity analysis was performed with the scvelo package (version 0.1.25)^33^. In order to match the original cell annotations, the matrices generated with the Alevin pipeline were filtered using the cell IDs that were kept in the initial analysis with RaceID algorithm. The filtered matrices were then used as input for the scvelo pipeline.

### Diffusion maps and diffusion pseudotime analyses

Diffusion maps and diffusion pseudotime were performed with the destiny package (version 2.14.0)^66^. Firstly, an object of the single cell experiment class was created with the SingleCellExperiment package (version 1.6.0) using as input the raw expression matrix obtained after filtering out the clusters corresponding to the INs, red blood cells, OPCs, pericytes, choroid plexus, microglia, Cajal-Retzius, subiculum and unknown clusters. The diffusion map was run with the DiffusionMap function using as values for the sigmas and k parameters the outputs of running the functions find_sigmas and find_dm_k, respectively. The diffusion pseudotime was subsequently computed using as input the diffusion maps object into the DPT function with default arguments.

### Cell fate bias estimation

Analyses of the cell fate bias probabilities were performed by applying a RNA velocity independent, FateID (version 01.9)^37^, and a RNA velocity dependent, CellRank (version 1.2.0)^36^, algorithm. Both analyses were run after filtering out the clusters corresponding to the INs, red blood cells, OPCs, pericytes, choroid plexus, microglia, Cajal-Retzius, and unknown clusters. Cell fate probabilities were computed after presetting CA1 and CA3 pyramidal, DG granule, Subic1 and Subic2 cell populations as the final states. The original tSNE projections computed by RaceID were provided as the dimensionally reduced coordinates for data visualization.

### Functional annotation analysis

To identify the main processes changing along the differentiation trajectory, functional over-representation analysis of gene ontology biological processes was performed with the R package clusterProfiler (version 3.12.0)^67^ using as input the genes represented in the respective nodes. For identification of the main biological processes related to the DEGs between control and cKOs in each of the individual clusters genes increased and decreased in expression were considered independently. Analyses of the main processes represented by the DEGs increasing in expression between the more mature and the less mature CA cell states, as well as in the case of the processes represented by the DEGs increasing or decreasing in expression among control and cKOs in the less mature CA cell state, were performed selecting the gene ontology for molecular functions. In all the independent analyses a significance *q-value* cutoff of 0.05 was selected.

### Transcription factors analysis

The exploratory analyses of the transcription factors changing in expression upon DOT1L depletion and differentially expressed among CA maturation states were performed by intersection of the respective DEGs with a list of *mus musculus* reported transcription factors obtained from the TRRUST (version 2) database (available at: https://www.grnpedia.org/trrust/), the database of transcription co-factors and transcription factor interactions (TcoF-DB, version 2) (available at: https://tools.sschmeier.com/tcof/doc/), and the AnimalTFDB (version 3.0) database (available at: http://bioinfo.life.hust.edu.cn/AnimalTFDB/#!/download).

### Analysis of the H3K79me2 profiles

The H3K79me2 ChIPseq data sets for NPC treated with DMSO or EPZ for 48h have been previously published^47^ and the files are accessible at the Gene Expression Omnibus database under the accession number GSE135318 (GSM4005219: NPC48h_DMSO_H3K79me2_rep2; GSM4005235: NPC48h_EPZ_H3K79me2_rep2). H3K79me2 profiles for the selected genes were plotted using the pyGenomeTracks package (version 3.6)^68^ and assembled in inkscape.

### Data accessibility, interoperability and reproducibility

The data files derived from the scRNAseq experiment discussed in this publication have been deposited in NCBI’s Gene Expression Omnibus^69^ and are accessible through GEO Series accession number GSE178105 (https://www.ncbi.nlm.nih.gov/geo/query/acc.cgi? acc=GSE178105). All bioinformatics analyses steps and settings are deposited under the link https://github.com/adsalas/DOT1L_hippocampus_development.

## Acknowledgments

ASB, MT, TV, HB, DG are members of the Deutsche Forschungsgemeinschaft (DFG, German Research Foundation)–funded 322977937/GRK2344 and thank for the support. DG was also supported by the Max Planck Society and the DFG (GR4980/3-1). The authors want to thank Dr. Mehmet Tekman (Department of Computer Science, Bioinformatics Group, University Freiburg) for providing suggestions regarding data analysis, and Dr. Sagar (Department of Medicine II, University Medical Center Freiburg) for his kind support during the libraries preparation for the scRNAseq experiment. Moreover, the authors thank Dr. Alejandro Villarreal, Dr. Henriette Franz, Dr. Nicole Hellbach, Stefanie Heidrich, Ute Baur (Institute of Anatomy and Cell Biology, Department of Molecular Embryology, University Freiburg) and HaiQin Zhang (RISE-Fellow) for performing some initial staining experiments and for help with animal experimentation. Further, the authors thank the Freiburg Galaxy Server team and all members of the Vogel group for discussion and their support.

## Author contributions

TV conceptualised the study. ASB, MT, HB and TV designed the experimental strategy. ASB performed the experiments. ASB and TV analyzed and interpreted data and results. JSH collaborated with the execution of the scRNAseq experiment and provided input for data analysis. DK contributed with smFISH experiments. MT analyzed data and prepared figures. MBS performed the quantification of exonic and intronic reads, and the scvelo analysis. HB, and DG supervised and provided input for the data analysis. ASB and TV wrote the manuscript and prepared the figures, with input from all co-authors. The manuscript was read and approved by all co-authors.

