## Supplementary_Figures for "Single-cell transcriptomic resolution of stem cells and their developmental trajectories in the hippocampus reveals epigenetic control of cell state perseverance"

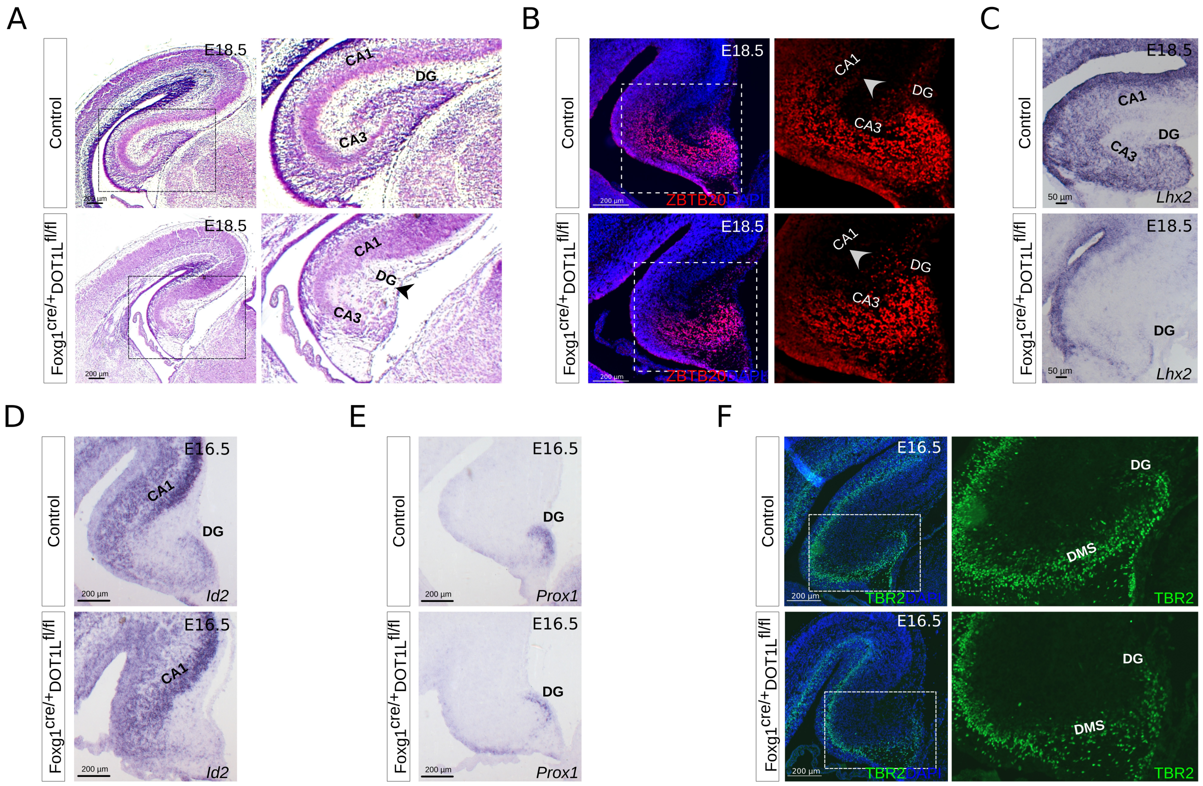


**Supplementary Figure 1.** A) Nissl staining of control and *Foxg1-Dot1l* cKO brains at E18.5 showing cytoarchitectural changes in the hippocampal area. cKO brains show thickening of the CA1 field and apparent loss of the DG (arrowheads) (n =3). B) ZBTB20 staining at E18.5 showing depletion of positive cells in the putative CA1 field upon DOT1L depletion (n =3). C) *Lhx2* ISH showing decreased expression at the level of the hippocampal region in the DOT1L mutants (n =3). A drastic decrease in *Lhx2* expression is seen in the intermediate zone and pyramidal cell layer as well as in the DG. D) Increased expression of *Id2* in the *Foxg1-Dot1l* putative CA1 field indicates ventro-lateral expansion of the subiculum (n =3). E) *Prox1* expression domain seems decreased at E16.5 upon DOT1L depletion (n =3). F) Less TBR2 positive cells migrate through the DMS and reach the DG in DOT1L mutants compared to controls at E16.5 (n =3). Scale bars are shown inside the image panels. CA: cornu ammonis; DMS: dentate migratory stream; DG: dentate gyrus.

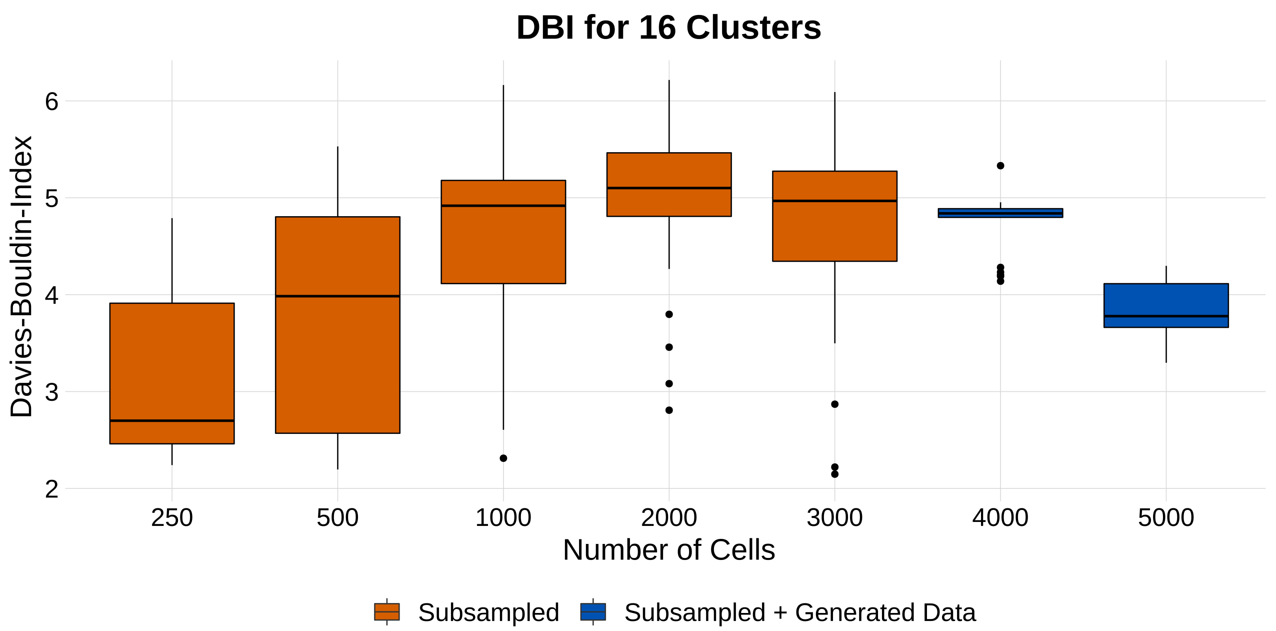


**Supplementary Figure 2.** Davies-Bouldin index indicating the quality of clustering for subsampled (orange) and subsampled in combination with simulated cells (blue) for pilot data of varying size. For each sample size, 30 random samples were taken followed by dimensionality reduction and clustering. Clusterings with less than 4000 cells exhibit high variability which indicates that sample size might be too small for reliably detecting cell identities.

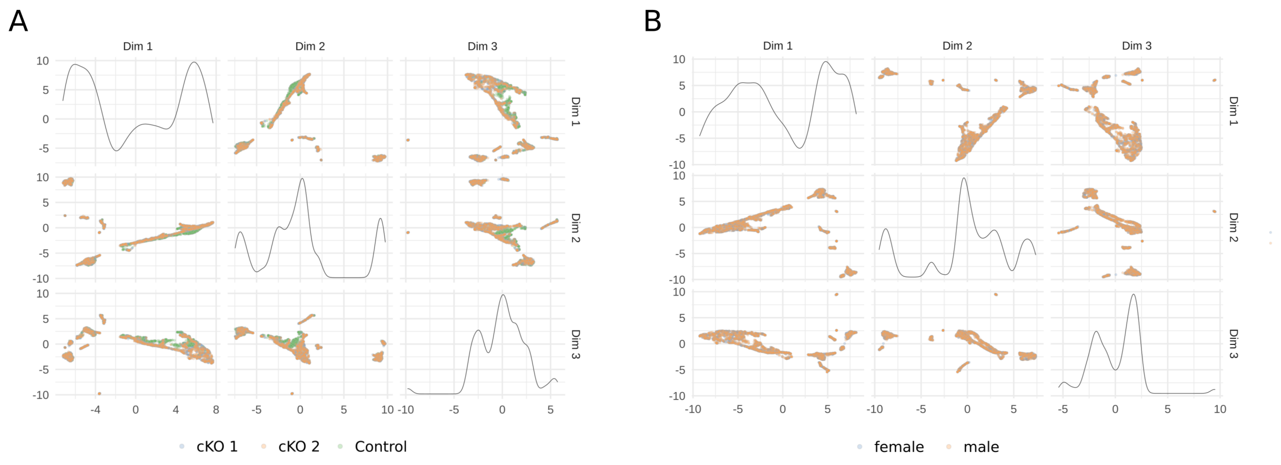


**Supplementary Figure 3.** Visual inspection of the effects of A) genotype and B) sex using denoised principal components analysis. A) The first three principal components coloured by the animals genotype. As was to be expected, slight differences between the genotypes are evident. B) First three principal components coloured by sex (comparing the 2 cKO animals). The visual analysis suggests no substantial differences between samples from distinct sex.

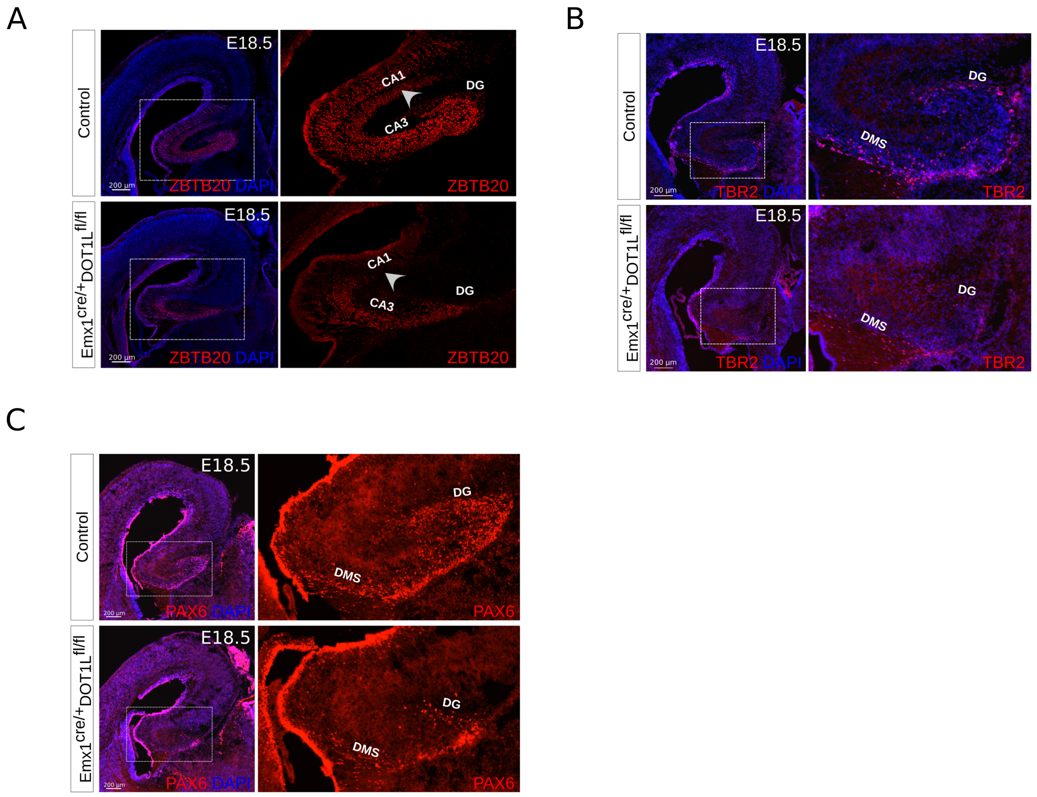


**Supplementary Figure 4.** A) ZBTB20 staining at E18.5 showing absence of positive cells in the putative CA1 field upon DOT1L depletion (arrowheads) (n =3). B) Less TBR2 positive cells migrate through the DMS and reach the DG in *Emx1-Dot1l* mutants compared to controls at E18.5 (n =3). C) Less PAX6 positive progenitor cells migrate through the DMS and reach the DG in *Emx1-Dot1l* mutants compared to controls at E18.5 (n =3). Scale bars are shown inside the image panels. CA: cornu ammonis; DMS: dentate migratory stream; DG: dentate gyrus.

A B

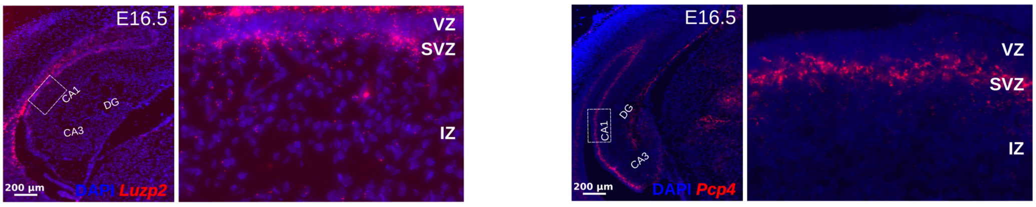


**Supplementary Figure 5.** smFISH stainings at E16.5 showing expression of A) *Luzp2* and B) *Pcp4*, both selected as a markers for the CA1 pyramidal cells belonging to the less mature cell state (n =3). Expression of both markers is higher at the level of the SVZ. Scale bars are shown inside the image panels. VZ: ventricular zone; SVZ: subventricular zone; IZ: intermediate zone.
