## Supplementary_Tables for "Single-cell transcriptomic resolution of stem cells and their developmental trajectories in the hippocampus reveals epigenetic control of cell state perseverance"

**Table 2.** Antibodies and dilutions used for immunofluorescence stainings.

| **Antibody** | **Species** | **Company / Catalog number** | **Dilution** |
| --- | --- | --- | --- |
| anti-PROX1 | Rabbit | Biolegend / 925201 | 1:1000 |
| anti-TBR2 | Rabbit | Abcam / ab23345 | 1:300 |
| anti-TBR2 | Mouse | Invitrogen /14487582 (Dan11mag) | 1:500 |
| anti-PAX6 | Rabbit | Biolegend / 901301 | 1:200 |
| anti-LHX2 | Rabbit | Abcam / ab184337 (EPR20449) | 1:200 |
| anti-SATB2 | Mouse | Abcam / ab51502 (SATBA4B10) | 1:200 |
| anti-NFIB | Rabbit | Sigma-Aldrich / HPA003956 | 1:200 |
| Anti-ZBTB20 | Rabbit | Sigma-Aldrich / HPA016815 | 1:200 |

**Table 3.** Probes and fluorophore dilutions used for single molecule FISH stainings.

| **Probe** | **Species** | **Catalog number** | **Dilution** |
| --- | --- | --- | --- |
| RNAscope®Probe -Mm-Hes5-C2 | Mouse | 400991-C2 | 1:800 |
| RNAscope®Probe -Mm-Sned1 | Mouse | 418571 | 1:800 |
| RNAscope®Probe -Mm-Wnt8b-C2 | Mouse | 405071-C2 | 1:800 |
| RNAscope®Probe -Mm-Sox21-C2 | Mouse | 429681-C2 | 1:800 |
| RNAscope®Probe -Mm-Adamts19 | Mouse | 500981 | 1:800 |
| RNAscope®Probe -Mm-Cybrd1-C3 | Mouse | 883131-C3 | 1:800 |
| RNAscope® Probe- Mm-Luzp2-C3 | Mouse | 492551-C3 | 1:1000 |
| RNAscope® Probe- Mm-Grp | Mouse | 317861 | 1:800 |
| RNAscope® Probe- Mm-Grik4-C2 | Mouse | 442021-C2 | 1:800 |
| RNAscope® Probe- Mm-Cpne4-C3 | Mouse | 474721-C3 | 1:800 |
| RNAscope® Probe- Mm-Tac2-C3 | Mouse | 446391-C3 | 1:800 |
| RNAscope® Probe- Mm-Pcp4 | Mouse | 402311 | 1:1000 |
| RNAscope® Probe- Mm-Pou3f1-C2 | Mouse | 436421-C2 | 1:800 |
